# Genomic basis of adaptation to serpentine soil in two *Alyssum* species shows convergence with *Arabidopsis* across 20 million years of divergence

**DOI:** 10.1101/2025.02.27.640498

**Authors:** Sonia Celestini, Miloš Duchoslav, Mahnaz Nezamivand-Chegini, Jörn Gerchen, Gabriela Šrámková, Raúl Wijfjes, Anna Krejčová, Nevena Kuzmanović, Stanislav Španiel, Korbinian Schneeberger, Levi Yant, Filip Kolář

## Abstract

**Background and Aims:** Serpentine outcrops, characterized by low nutrient availability, high heavy metal concentrations, propensity to drought, and island-like distributions, offer valuable systems to study parallelisms in repeated adaptation to extreme environments. While shared phenotypic manifestation of adaptation to serpentine environments has been investigated in many species, it is still unclear whether there may be a common genetic basis underlying such responses. Here we assess local adaptation to serpentine soil and infer the parallel genetic signatures of local adaptation to serpentine environments in two thus far unexplored closely related species, *Alyssum gmelinii* and *Alyssum spruneri* (Brassicaceae). Then we measure gene- and function-level convergence with the previously explored *Arabidopsis arenosa*, to reveal candidate shared adaptive strategies within Brassicaceae.

**Methods:** We tested for adaptation using a reciprocal substrate-transplant experiment in *A. gmelinii*. Then, after assembling a reference genome, we generated population-level sequencing data of four population pairs and performed genome scans for directional selection to infer serpentine adaptive candidate genes in *Alyssum*. Finally, we compared candidate gene lists with those inferred in similar experiments in *Arabidopsis arenosa* and used protein-protein interaction networks to discern functional convergence in serpentine adaptation.

**Key Results:** Independent colonization of serpentine environments by *Alyssum* populations is associated with footprints of selection on genes related to ion transport and homeostasis, nutrient and water uptake, and life-history traits related to germination and reproduction. Reciprocal transplant experiments demonstrated that adapted plants germinate sooner and exhibit better growth in serpentine conditions while excluding heavy metals and increasing Ca uptake in their tissues. Finally, a significant fraction of such genes and molecular pathways is shared with *Arabidopsis arenosa*.

**Conclusions:** We show that genetic adaptation to the multi-factorial challenge imposed by serpentine environments involves key pathways that are shared not only between closely related species, but also between Brassicaceae tribes of ∼20 Mya divergence.

## Introduction

The remarkable ability of plants to adapt to diverse hostile environments has long fascinated evolutionary biologists. Serpentine barrens offer powerful models to address this topic (Konečná *et al*., 2020). Derived from the weathering of ultramafic rocks, serpentine soils present variable and multi-factorial challenges, summarised as “the serpentine syndrome” (Jenny, 1980). This illustrates the cumulative effect of the three main stressful components - chemical, physical and biotic - on plant growth and survival. Despite considerable site-to-site variation in the content of particular elements, serpentine soils are principally defined by a highly skewed Ca/Mg ratio, with greatly reduced availability of Ca and increased availability of Mg (Harrison and Rajakaruna, 2011). Other characteristics are the low concentration of important macronutrients alongside the high content in heavy metals such as Co, Cr and Ni (Proctor and Woodell, 1975; Brady *et al*., 2005; Harrison and Rajakaruna, 2011). Physical stress on plant colonists comes from serpentines’ porous texture and dark colour, translating into frequent droughts and soil overheating. Biologically, serpentine sites represent islands of abrupt change in ecosystem productivity, resulting in low competition and high rates of endemism (Kruckeberg, 1984; Harrison, 1999a; b; Anacker, 2014). Taken together, these challenging conditions impose strong selective pressures (Brady *et al*., 2005; Konečná *et al*., 2020), making adapted populations valuable model systems to study plant tolerance strategies to environmental hazards.

Long of interest to ecologists and plant biologists, the phenotypic manifestations of adaptations to serpentine soil have been extensively characterised. Solutions often involve differential ion uptake and homeostasis, exclusion or compartmentalization of heavy metals and shifts in life-history traits (Berglund and Westerbergh, 2001; Wright *et al*., 2006; Sambatti and Rice, 2006; Palm *et al*., 2012; Kolář *et al*., 2014; Chathuranga *et al*., 2015; Veatch-Blohm *et al*., 2017; Salehi Eskandari *et al*., 2017; Mišljenović *et al*., 2020; Lazzaro *et al*., 2021; Dittmar and Schemske, 2023; Bürki *et al*., 2024). Yet, these works have been only rarely complemented by investigation of the underlying adaptive genetic basis, mostly focusing on the *Arabidopsis* genus (Turner *et al*., 2008, 2010; Arnold *et al*., 2016; Sobczyk *et al*., 2017; Selby and Willis, 2018; Konečná *et al*., 2021). Additionally, the island-like distribution of serpentines (Roberts and Proctor, 1992) is of particular interest for evolutionary biology as it often triggers parallel adaptation in each serpentine site independently (Rajakaruna *et al*., 2003; Mengoni *et al*., 2003; Berglund *et al*., 2004; Sakaguchi *et al*., 2019; Lazzaro *et al*., 2021; Konečná *et al*., 2021). These natural replicates of adaptation can be leveraged to discern consistent trends in mechanisms and possibly genetic basis of local substrate adaptation (Rajakaruna, 2004; Konečná *et al*., 2020), which helps predicting evolutionary outcomes (Bohutínská and Peichel, 2024). As a proof-of-concept, the genomic studies provided strong evidence for shared genetic basis of serpentine adaptation among populations within a species (Turner *et al*., 2010; Selby and Willis, 2018; Konečná *et al*., 2021) and between sister species (Arnold *et al*., 2016). However, it remains unknown whether there is also a convergent genetic basis over broader evolutionary timescales, across genera and families (Konečná *et al*., 2020).

Here we investigate genetic adaptation to serpentine conditions in two so far unexplored Brassicaceae species which often occur on and off serpentine soil: the closely related species *Alyssum gmelinii* and *Alyssum spruneri* from the *Alyssum montanum-A. repens* group (Španiel *et al*., 2011, 2012; Španiel, Zozomová-Lihová, *et al*., 2017; Španiel, Marhold, *et al*., 2017). This complex is notorious not only for its challenging taxonomy, shaped by recent rapid diversification in the early Pleistocene and multiple polyploidization events (Cetlová *et al*., 2021; Španiel *et al*., 2023), but also for its ecological diversity. *Alyssum gmelinii* can be found on sandy dunes as well as on rocky outcrops on various substrates in eastern and central Europe. *Alyssum spruneri* grows in the central Balkans on rocky outcrops and both species are known to occupy serpentine environments. Both species are perennial outcrossers, with limited pollen and seed dispersal, and both are present as both diploid and polyploid cytotypes (Španiel *et al*., 2011; Španiel, Zozomová-Lihová, *et al*., 2017; Španiel, Marhold, *et al*., 2017). Belonging to the *Alysseae* tribe, they share a close phylogenetic history with *Odontarrhena*, a well-studied genus in terms of serpentine adaptation and metal tolerance, that taxonomists recently separated from *Alyssum* (Warwick *et al*., 2008; Cecchi *et al*., 2010; Rešetnik *et al*., 2013; Španiel *et al*., 2015; Li *et al*., 2015). Notably, *Odontarrhena* includes the greatest number of nickel-hyperaccumulating species in the plant kingdom (Brooks *et al*., 1979), which have garnered considerable attention for their phytoremediation potential (Li *et al*., 2003; Broadhurst *et al*., 2004; Seneviratne *et al*., 2016; Sobczyk *et al*., 2017; Nkrumah *et al*., 2021). However, most plants adapted to ultramafic soils are not hyperaccumulators. This highlights the need to study non-hyperaccumulating species such as those within *Alyssum*, which can provide broader insights into the mechanisms underlying plant tolerance to serpentine conditions. Moreover, we can directly compare our findings with the previous body of research in *Arabidopsis* to explore patterns of convergence.

We combine transplant experiments and whole genome sequencing of serpentine–non-serpentine population pairs in order to infer the repeated genomic basis of serpentine adaptation both within *Alyssum* and also more broadly in Brassicaceae. We specifically ask: i) Do serpentine-endemic populations of *A. gmelinii* show a fitness advantage on serpentine soil compared to non-serpentine populations? ii) Is there a shared genetic basis underlying serpentine adaptation between the two closely related *Alyssum* species? iii) Do the candidate adaptive genes, or their inferred functions, detected in *Alyssum* exhibit footprints of convergent selection with the distantly related *Arabidopsis arenosa*?

## Materials and Methods

### Sampling

To perform a robust set of divergence selection scans and thus determine candidate genetic bases of serpentine adaptation in the *Alyssum montanum-A. repens* complex, we sampled a set of four spatially proximate serpentine–non-serpentine (S-N) population pairs. Based on close study of the species distribution (Španiel *et al*., 2011; Španiel, Marhold, *et al*., 2017; Španiel *et al*., 2023) we identified four pairs of geographically close S-N populations of *Alyssum gmelinii* and *Alyssum spruneri* (two pairs per species) in Central and Southern Europe. To generalise our findings, we included populations of two ploidy levels for each species: diploid and autotetraploid. Distance between population pairs ranged from 4 km to 26 km, and each pair closely coincided in elevation (Supplementary Data S1, Fig. 1A).

**Fig. 1.**
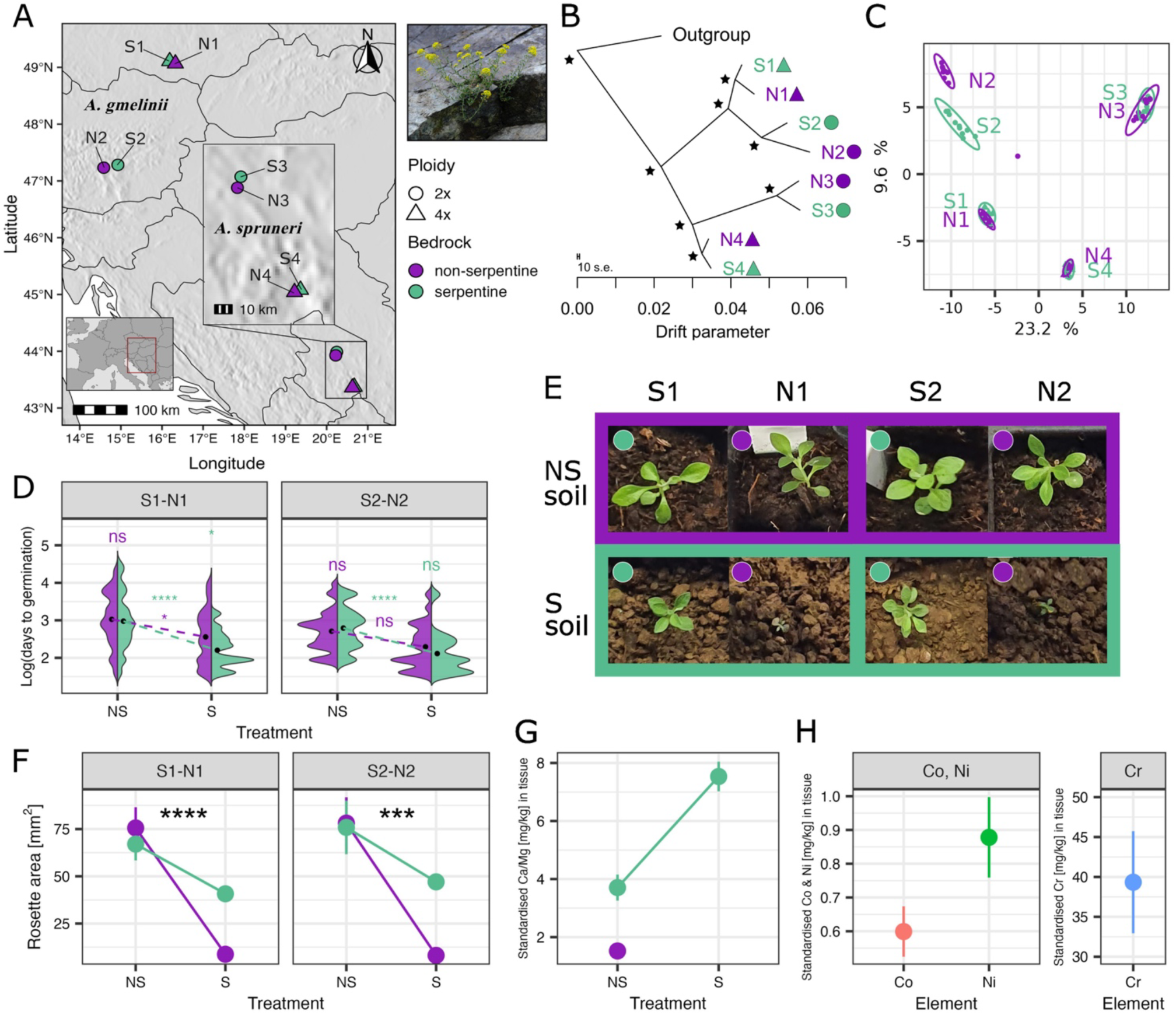
Repeated colonization and adaptation of the two *Alyssum* species to serpentine soil. (A) Geographical distribution of the investigated populations in Central and Southeastern Europe. Each population pair, represented with a specific number (1-4), is composed of a serpentine (S, green) and a non-serpentine (N, violet) population which are geographically proximate, symbols depict different ploidy levels within each species (diploid and autotetraploid). On the right, illustrative photo of an individual from population S2 growing on a serpentine rock (photo by S. Celestini). (B) Allele frequency covariance graph showing relationships among the studied populations calculated using 1,082,883 neutral fourfold-degenerate SNPs (4dg) in Treemix, asterisk indicate the 100% bootstrap branch support. The outgroup represents a diploid *Alyssum repens* population from Austria. (C) Principal Component Analysis (PCA) of the individual genotypes calculated from 10,000 4dg SNPs. Individuals cluster by population of origin, while PC1 axis splits populations by species and PC2 by distinct region, also corresponding with ploidy. (D) Proportion of newly germinated seeds per day after sowing (log-transformed) for each population pair used in the transplant experiment, showcased through mirrored density plot and means (black dots). (E) Photos illustrating adaptive response to serpentine soils (green frame) in the two population pairs 1-2 depending on the soil of origin (dot colour) (photos by S. Celestini). (F) Differences in rosette area in the population pairs 1-2. Significance of the soil treatment by soil of origin interaction in the linear mixed effect model is indicated by asterisks for each pair: β = 1.503, SE = 0.300, t = 5.001, p = 2.71e-06 for pair 1; β = 1.475, SE = 0.270, t = 5.449, p = 3.28e-07 for pair 2). (G) Differences in Ca/Mg ratio [mg/kg] uptake in root and shoot tissues of plants cultivated in S and NS soils, standardised by the amount of Ca/Mg [mg/kg] ratio present in the treatment soil. (H) Levels of cobalt (Co), nickel (Ni) and chromium (Cr) uptake in root and shoot tissues of plants of serpentine origin cultivated in serpentine conditions, standardised by their values in the cultivation soil. Points denote mean, error bars depict standard error of mean in figures F, G and H. Colours (green and violet) refer to the soil type of origin (S and N respectively) throughout the figure.

We sampled and silica-dried leaf material for ten individuals per population for genome sequencing and fresh tissue for confirmation of their ploidy level by means of flow cytometry following our standard two-step protocol with Otto buffers and DAPI staining as in Kolář *et al*. (2016). From populations of *A. gmelinii*, we also sampled seeds from a minimum of 15 mother plants. To get an outgroup for some analyses, we also sampled 10 individuals from a diploid population of *Alyssum repens* from Austria (see Supplementary Data S1 for locality details). We gathered information about soil condition at each location collecting ∼ 1 litre of topsoil at three different spots spanning the population area. Soil samples from each population were mixed, air-dried, sieved and analysed at the Analytical Laboratory of the Institute of Botany, Průhonice, in Czech Republic (see Supplementary Methods, Supplementary Data S1). Collection of Austrian populations was granted by the Styrian Provincial Government, Department of Nature and General Environmental Protection (permission No. ABT13-151190/2021-7).

### Reference Genome Sequencing and Assembly

We generated a de novo genome assembly using long DNA reads sequenced using the PacBio Revio platform from one diploid individual of *Alyssum gmelinii* sampled in population S2.

A total of 0.2 g leaf material from one individual plant was ground using liquid nitrogen and isolated as described in Russo *et al*. (2022). The quantity and quality of high molecular weight DNA was checked on a Qubit Fluorometer 2.0 (Invitrogen) using the Qubit dsDNA HS Assay kit and the NanoDrop One spectrophotometer (Thermo Scientific). Fragment sizes were assessed using the Genomic DNA Tapestation assay (Agilent).

Sequence libraries were constructed using the PacBio HiFi long-read single molecule real-time (SMRT) sequencing according to the manufacturer’s instructions (PacBio, Menlo Park, CA, USA). Library preparation and PacBio sequencing was performed at Novogene Europe on a Revio platform. A total of 2,344,090 HiFi consensus reads (average length: 16.0 kbp) resulted in ∼37.4 Gbp of sequence data.

To characterize the reference genome we first performed k-mer analysis on the sequencing data using KMC (Kokot *et al*., 2017), followed by genome size and heterozygosity estimation with GenomeScope2, using K=31 (Ranallo-Benavidez *et al*., 2020). These analyses provided initial insights into genome complexity and guided the sequencing depth requirements for the assembly. Hifiasm-0.19.8 (Cheng *et al*., 2021) was then employed for *de novo* assembly of the PacBio HiFi reads, using the default settings, except for the –hom-cov 75 parameter. Three different methods were used for quality assessment of the assembly; QUAST v.5.2.0 (Gurevich *et al*., 2013) provided assembly statistics, BUSCO v.5.4.3 (Simão *et al*., 2015) was used to assess genome completeness with the brassicaceae_odb10 lineage dataset, and Merqury (Rhie *et al*., 2020) provided a quality value (QV) score indicating the precision of the base calls and assessed the assembly completeness. We used Meryl v.1.3 (Rhie *et al*., 2020) to count k-mers and construct the Meryl database required for Merqury.

To refine the assembly and minimize redundancy, we used purge_haplotigs v.1.1.3 (Roach *et al*., 2018). This step allowed us to identify and filter out haplotigs, resulting in a more accurate representation of the primary haplotype by alterate haplotypes in our pseudohaploid assembly. Finally, we identified and removed non-nuclear DNA by blasting the primary assembly against a reference plastid sequence database from RefSeq (O’Leary *et al*., 2016) using MMseqs2 v.2.14 (Steinegger and Söding, 2017) to detect and filter plastid DNA, including mitochondrial and chloroplast sequences. Repeat elements were identified using RepeatModeler v.1.4 (Flynn *et al*., 2020), which generated a custom repeat library for *Alyssum gmelinii*. This library was then used with RepeatMasker v.4.1.2 (Tarailo-Graovac and Chen, 2009) to annotate repetitive elements in the assembly.

### Gene prediction, annotation and orthology detection

We first trimmed RNA-seq data to remove sequencing adapters and low-quality regions using Trim Galore v.0.6.2 (https://github.com/FelixKrueger/TrimGalore) with paired mode (‘--paired’) and aligned to the assembled reference genome using HISAT2 v.2.1.0 (Kim *et al*., 2019) with command line options ‘-q -t --dta’. The resulting SAM files were converted to BAM files, sorted and merged using SAMtools v.1.14 (Li *et al*., 2009).

We predicted the protein-coding genes by BRAKER v.3.0.8 (Gabriel *et al*., 2024) with default settings based on our RNA-seq data and on Viridiplantae proteins from OrthoDB v.11 (Kuznetsov *et al*., 2023). To include regions that are transcribed but do not encode proteins (non-coding RNA genes and similar features), we filtered transcripts assembled from RNA-seq data using StringTie v.2.2.3 (Pertea *et al*., 2015) with default settings using bedtools v.2.30.0 (Quinlan and Hall, 2010) to keep only those that did not have any overlap with protein-coding gene annotations. We included them in the annotation as ‘ncRNA_gene’ and ‘ncRNA’. We converted GTF to GFF using AGAT v.1.4.0 (Dainat *et al*., 2024). We changed feature IDs using a custom R script.

To assign *Arabidopsis thaliana* orthologues to *Alyssum* genes, we run OrthoFinder v.2.5.4 (Emms and Kelly, 2019) with proteomes of 18 Brassicaceae species with reference genomes and annotations of suitable quality as input (*Alyssum gmelinii, Arabidopsis arenosa, Arabidopsis lyrata, A. thaliana, Arabis alpina, Brassica oleracea, Brassica rapa, Camelina sativa, Capsella rubella, Cardamine glauca, Cardamine hirsuta, Cochlearia excelsa, Conringia planisiliqua, Euclidium syriacum, Eutrema salsugineum, Noccaea praecox, Raphanus sativus, Rorippa islandica*). The resulting *Alyssum - A. thaliana* orthologues were further supplemented by best scoring hits from BLAST 2.16.0+ (Camacho *et al*., 2009) search of *Alyssum* protein sequences against *A. thaliana* protein sequences. Only hits with percentage of identical matches > 40 and query coverage per local alignment > 50% were considered. If several *A. thaliana* genes were assigned as orthologues to one *Alyssum* gene by OrthoFinder, the best scoring in BLAST search was selected.

The details of genome annotation and orthology detection and corresponding scripts are available at https://github.com/mduchoslav/Alyssum_serpentine_paper.

### DNA extraction and library preparation for short read sequencing

We extracted genomic DNA from silica-gel-dried leaves using the sorbitol extraction method and then purified it using AMPure XP (Beckman Coulter Inc., Brea, California, USA). Shotgun sequencing followed genome sequencing protocol LITE of Perez-Sepulveda *et al*. (2021). The libraries were sequenced in 300 cycles (2 × 150 bp paired-end/PE reads) on the Illumina NovaSeq platform.

### Variant calling and filtration

We conducted the analysis using a bioinformatics workflow implemented using Snakemake (Mölder *et al*., 2021) described at https://github.com/jgerchen/polyploid_variant_calling. This pipeline was designed specifically for mixed-ploidy populations, generally following best-practices (Bohutínská *et al*., 2023), and includes steps for quality control, read alignment, variant calling, and filtration.

Specifically, we conducted initial quality control of raw sequencing data using fastqc v.0.12.1 (Andrews, 2010), then we trimmed and filtered reads of low quality and adaptors using trimmomatic v.0.39 (Bolger *et al*., 2014) with default parameters. We mapped the refined reads using bwa mem v.0.7.18 (Li and Durbin, 2009) with default settings. Duplicate reads were marked using Picard tools v.3.1.1 (http://broadinstitute.github.io/picard). Afterwards, we called genotypes per sample using the HaplotypeCaller module of GATK v.4.6.0.0 (Poplin *et al*., 2017), specifying the ploidy of each individual using the --ploidy argument and allowing for a minimum base quality score of 20, a minimum mapping quality of 20, and a maximum genotype count of 350. We then combined samples in a GenomicsDB using the GenomicsDBImport module and performed combined genotyping of all populations using the GenotypeGVCFs module with genotyping of invariant sites activated. Filtration of variants was performed using bcftools v.1.20 (Danecek *et al*., 2021). We first split the unfiltered dataset into variants and invariant sites. We filtered variant sites to retain only biallelic SNPs and kept SNPs that passed the GATK-recommended ‘best practices’ filtering parameters (QD>2, FS<60, MQ>40, MQRankSum>-12.5, ReadPosRankSum>-8 and SOR<3). Then, we set individual genotypes that had a sequencing depth smaller than 9 to no-call and removed sites with more than 50% missing data. We also filtered genotypes from the subset of invariant sites and set genotypes with a sequencing depth smaller than 9 to no-call and removed sites with more than 50% missing data. We merged the filtered datasets for biallelic and invariant sites and applied a depth mask filter using a custom python script that removed sites where more than half of individuals had a sequencing depth greater than two standard deviations above the average depth per sample. We identified fourfold degenerate sites from the GFF annotation file using degenotate pipeline (https://github.com/harvardinformatics/degenotate?tab=readme-ov-file#degeneracy-per-site-bed-file) and then extracted. We further filtered based on depth and proportion of missing data specifically for particular analyses (detailed below).

### Site frequency spectra

We calculated unfolded site frequency spectra from population allele frequencies using a custom python script (https://github.com/jgerchen/polyintro/tree/main/workflow/scripts/SFS), which accounts for missing data in one individual by subsampling alleles to a depth of 18 for diploid populations and a depth of 36 for tetraploid populations, while discarding variants with more missing data. The input file included all bi-allelic SNPs (25,458,120 SNPs, with an average depth of 33x and 7% missing data) and invariant sites.

### Population structure analysis

To investigate population genetic structure, relatedness, and genetic diversity, we analysed putatively neutral fourfold degenerate (4dg) biallelic SNPs with a maximum individual genotype missingness of 0.5 and a minimum depth of 8x, yielding 1,082,883 SNPs with an average depth of 32x and 4.5% missing data. We first displayed relationships among individuals using principal component analysis (PCA) calculated on individual genotypes using glPcaFast function (https://github.com/thibautjombart/adegenet/pull/150) in R, focusing on the first 10,000 SNPs. To further explore relationships among populations, we constructed an allele frequency covariance graph using TreeMix v.1.13 (Pickrell & Pritchard, 2012). Input files for TreeMix were prepared with Python scripts from https://github.com/mbohutinska/TreeMix_input, filtering the VCF files to allow a maximum of 20% missing genotypes. We used the diploid *Alyssum repens* population as an outgroup. Tree topology was tested with bootstrapping (block size of 1 kb, matching the selection scan window size; see below) across 100 replicates, and results were visualized using FigTree v.1.4.4. We assessed model fit by running the analysis with 0 to 6 migration edges to capture potential gene flow between populations. For within-population diversity metrics, we calculated nucleotide diversity (π) and Tajima’s D (Tajima, 1989), down-sampling to seven individuals per population to account for missing data and reduce sample size bias. These analyses were conducted with custom Python scripts from the ScanTools toolkit (https://github.com/sonia23cld/ScanTools_ProtEvol) with a 50,000-bp window size and a minimum of 10 SNPs per window. Additionally, we estimated population differentiation using Rho, an alternative to F_ST_ suited for comparisons across ploidy levels (Ronfort et al., 1998; Meirmans et al., 2018), for each pair of populations. Finally, we used Entropy v.2.0 (Shastry *et al*., 2021), a hierarchical Bayesian method adapted for mixed-ploidy datasets, to infer population clustering and ancestry. Input data consisted of pruned and filtered 4dg VCF files (6,686 SNPs, with 3.09% missing data). We initialized admixture estimates by applying k-means clustering to genotype likelihoods obtained from linear discriminant analysis in the MASS package v.7.3-57 (Venables WN and Ripley BD, 2002) in R, to support Markov chain Monte Carlo (MCMC) convergence. Entropy was then run in parallel (Tange, 2011) with K values from 2 to 5, using 60,000 MCMC steps, discarding the first 20,000 as burn-in, and thinning every 20 steps. The resulting genotype estimates were visualized in R (https://github.com/sonia23cld/Entropy).

### Reciprocal transplant experiment

To investigate adaptation to serpentine conditions in our study group, we designed a transplant experiment to compare fitness traits of plants reciprocally grown in serpentine and non-serpentine soils. To focus on the effects of soil type, we cultivated the plants under uniform and controlled greenhouse conditions. Due to logistical constraints, our study focused on the four Central European populations of *A. gmelinii* (population pairs 1 and 2) previously described. We sowed 10 seeds per maternal plant from 10 mothers per population (except for N2, where we used additional maternal plants to achieve a similar seed count), resulting in a total of 720 seeds. To keep standard conditions across the experiment, we used commercial garden soil as the control, and natural serpentine soil, collected from the “Borovsko” locality near Prague, Czech Republic (Latitude: 49.683; Longitude: 15.133), as the treatment soil for the experiment. Although this serpentine locality differs geographically from the origin of the populations used in this study, its soil elemental composition is similar to the original *A. gmelinii* localities (Supplementary fig. S1). Seeds were germinated in Petri dishes in control and treatment soil under standard growth chamber conditions (daytime temperature 23°C, nighttime 12°C). Three weeks after sowing, we transplanted approximately three seedlings per maternal plant per treatment into individual pots in multipot trays, totalling 256 seedlings, and moved them to the greenhouse. Our objective was to assess the fitness effects by examining the interaction between soil treatment and soil of origin, using proxies such as germination time, rosette area, final biomass, and elemental accumulation in plant tissues. Plants were left growing in the greenhouse for an additional 67 days, during which they were watered and rotated twice a week until they reached a vegetative growth plateau. At the end of this period, we harvested the entire plant biomass (above and below ground), rinsed it with distilled water, oven-dried at 75°C for two nights and weighted it. Elemental analysis of the dried tissue for Ca, Mg, Co, Cr, and Ni in plant tissues was then conducted at the Institute of Environmental and Chemical Engineering, Faculty of Chemical Technology, University of Pardubice, using inductively coupled plasma optical emission spectrometry (ICP-OES; see Supplementary Methods). Given the high mortality/small biomass attained by the non-serpentine origin plants in serpentine treatment, we lack ionomic data relative to these populations (N1, N2) grown in serpentine treatment.

Statistical analysis of the collected traits was performed in R v.4.4.2. Differences in *germination rate* were tested fitting a generalized linear mixed-effects model (GLMM) using the binomial distribution. The response variable was the number of germinated seeds out of the total seeds sown. Soil type of treatment, mother plant’s soil of origin, and their interaction, were included as fixed effects, while the maternal code was included as a random effect. The model was fitted using the glmer() function in the lme4 package (Bates *et al*., 2015). Differences in *germination time* were tested using survival analysis with a mixed-effects Cox proportional hazards model, implemented in the coxme v.2.2-22 package (Therneau, 2024). The survival outcome was days to germination, with censoring for seeds that did not germinate during the full experimental period. Fixed effects included soil treatment (factor: “S” and reference soil) and seed’s soil of origin (factor: “S” and reference origin), together with their interaction. Random intercepts were included for the population of origin and maternal code to account for population-level and maternal effects. Hazard ratios (HR) are reported to quantify the magnitude of effects. Difference in *plant rosette area* was tested at 25 days after repotting, i.e. before death rates of originally non-serpentine plants in serpentine treatment started to increase. We tested the effect of soil of origin, soil of treatment, and their interaction on the scored rosette area using linear mixed effects models with the function lmer from the r package lmerTest v.3.1-3, (Kuznetsova *et al*., 2017). Data was normalized using Box Cox Transformation implemented in the bestNormalize R package v.1.9.0 (Peterson and Cavanaugh, 2020) to meet model assumptions. Plants’ age, treated as a continuous covariate, was also included as a fixed effect to account for potential variability in plant size due to differences in their germination time. To account for non-independence among individuals with the same maternal origin, maternal identity was included as a random intercept. Model assumptions, including normality of residuals and homogeneity of variance, were visually inspected using diagnostic plots. Finally, we used the same model to test differences in the total *plant biomass* scored at the end of the experiment, normalizing the data using the Ordered Quantile (ORQ) normalization transformation implemented in the bestNormalize R package v.1.9.0 (Peterson and Cavanaugh, 2020). Here, however, we missed a large proportion of data for populations N1 and N2 cultivated in serpentine soil, as most of the plants died during the experiment.

### Window-based scans for directional selection

In order to detect genetic footprints of directional selection in response to serpentine, we designed three window-based approaches leveraging S-N population differentiation which conservatively allowed us to identify candidate genes for repeated adaptation to serpentine conditions in four population contrasts across two *Alyssum* species. For all the three analyses, we used as input a vcf file including all biallelic SNPs and variants filtered as explained in the previous section (25,458,120 SNPs, with an average depth of 33x and 7% missing data). We estimated appropriate window size by calculating genotypic correlation (r^2^), i.e. the proxy linkage-disequilibrium (LD) decay, for all diploid populations using PopLDdecay (Zhang *et al*., 2019). Then, we used ScanTools pipeline and calculated pairwise (S-N populations) Hudson F_ST_ (Hudson *et al*., 1992) for non-overlapping 1 kbp windows (LD was considered low based on Supplementary fig. S2) along the genome, excluding potential non-informative windows with less than 10 SNPs.

As a first method, we selected the 1% quantile windows showing the highest F_ST_ values in each population pair. Secondly, for each S-N pair we applied a modified version of Population Branch Statistics (PBS), Population Branch Excess (PBE), as defined in Yassin *et al*., (2016):

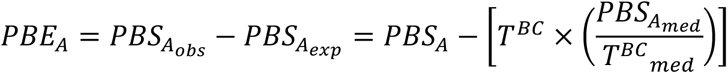

Here, the observed PBS is compared to an expected PBS and the resulting positive values will reflect positive selection specific to the focal branch. In our analysis, we used as focal branches (branch A) always the serpentine populations, while as nonfocal and closely related populations (branch B) we used the non-serpentine population from the same region (pair) as branch A. Finally, as a more distantly related outgroup nonfocal populations (branch C) we used the remaining non-serpentine population belonging to the same species. Specifically, for each focal (S) population we designed the analysis (branch A-B-C respectively) as follows: S1-N1-N2, S2-N2-N1, S3-N3-N4, S4-N4-N3. We then retained the 1% quantile outlier windows with the highest calculated PBE value. Then, we overlapped, within each pair, the results of these two methods obtaining four lists of “population pair-specific” candidate genes. In order to get a set of genes that would represent a consensus between the two species in our dataset, we then overlapped these four lists and retained the genes that were consistently outliers in at least two population pairs (referred to as “top candidates”). Complementarily, we analysed repeated patterns in the window-based F_ST_ values across the complete dataset using PicMin (Booker *et al*., 2023), a method that uses the theory of order statistics to quantify, for each window, the non-randomness of repeated molecular evolution across several population pairs (available at https://github.com/TBooker/PicMin). We ran PicMin with the ɑAdapt set to 0.05 (appropriate for a test of 4 lineages) and 10k repetitions. We then corrected the output p-values for false-discovery rate with FDR-correction (q) and kept only windows with q lower than 0.05 for further analysis. For each of the three methods, we then identified the genes (including 5’ untranslated regions (UTRs), start codons, exons, introns, stop codons, and 3’UTRs) overlapping the retained outlier windows. The statistical significance of the overlap between the gene lists resulting from each method was assessed by comparing it to the expected number of overlapping items given the sample size using Fisher’s exact test in SuperExactTest R package (Wang *et al*., 2015).

To infer the functional involvement of each list of candidate genes in potential biological processes, we used *A. thaliana* orthologs and ran Gene Ontology (GO) enrichment analysis using the R package clusterProfiler v.4.14.4 (Xu *et al*., 2024), with the function enrichGO() and TAIR gene identifier for gene ontology over-representation test. GO terms were annotated using the org.At.tair.db reference database, significance was assessed using hypergeometric test and p-values were adjusted for multiple testing using the Benjamini-Hochberg correction. Significantly enriched GO terms were selected based on p-value threshold of 0.05. We then visualised the enriched GO terms for each gene list using an heatmap from ComplexHeatmap v.2.22.0 R package (Gu, 2022).

### Assessing signal of potential adaptive convergence between species

We assessed the degree of genetic and/or functional serpentine soil adaptive convergence between *Alyssum* and the only other recently investigated Brassicaceae species *A. arenosa*. To do so, we used *A. arenosa* serpentine adaptation candidate genes detected in five S-N population pairs from (Konečná *et al*., 2021). To make our results comparable in terms of candidate gene selection criteria, we used the list of parallel candidate genes defined as top 1% F_ST_ windows outliers identified in at least two *A. arenosa* population pairs (222 orthologs *A. thaliana* genes). We first tested the extent of genetic overlap (in terms of *A. thaliana* orthologs) between species using Fisher’s exact test in the SuperExactTest R package (Wang *et al*., 2015). We compared *A. arenosa* candidate genes with *Alyssum* lists of top candidates and PicMin candidates separately. As background universe to compute the statistical significance of the observed intersections, we used the 19,908 *A. gmelinii* – *Arabidopsis* shared *A. thaliana* orthologs identified by OrthoFinder v.2.5.4 (Emms and Kelly, 2019). Similarly, we estimated the extent of adaptive functional overlap comparing the lists of significantly enriched GO-terms for each category. For this, we performed Gene Ontology (GO) enrichment analysis for *A. arenosa* candidate genes using the R package clusterProfiler v.4.14.4 (Xu *et al*., 2024) as explained above and tested for the significant overlap using Fisher’s exact test in SuperExactTest R package (Wang *et al*., 2015). Here the background population size was set as the maximum available number of “biological processes” GO terms for the *A. thaliana* orthologs in the package org.At.tair.db v.3.20 (Carlson, 2024).

Finally, we investigated potential convergence at the level of functional pathways by calculating the level of protein-protein interactions between candidate genes detected in each genus. We constructed a functional protein interaction network (PPIN) using nodes and edges downloaded from STRING v.12 (https://string-db.org/) (Szklarczyk *et al*., 2023) protein network database (scored links between proteins) for *A. thaliana* proteins. We associated each protein sequence Ids (AGI) to a UniProt accession using TAIR10 release of IDs conversions (https://www.arabidopsis.org/download/list?dir=Proteins%2FId_conversions) (Berardini *et al*., 2015). To build the network, we retained only the edges directly connecting *A. arenosa* nodes (218 proteins associated to *A. arenosa* candidate genes) to *A. gmelinii* nodes (127 proteins associated to consensus *Alyssum* candidate genes – here the union of the top candidates and the PicMin candidates), and vice-versa. This allowed us to focus only on the interactions between-species and not within, avoiding inflating the number of the edges. Note that 13 proteins overlapped between species. We then visualised specific portions or the PPIN in R using the packages igraph v.1.3.1 (Csardi and Nepusz, 2006) and tcltk v.4.1.2 (Grosjean, 2022). Functional enrichment of the network nodes was investigated running Gene Ontology (GO) enrichment analysis using the R package clusterProfiler v.4.14.4 (Xu et al., 2024) as above. Finally, we tested if the observed number of interactions (edges) was significantly higher than random performing a one-sided permutation test with 5,000 permutations in R, at each turn scoring the number of edges between *A. gmelinii* nodes and a randomly selected set of 205 *A. thaliana* proteins.

## Results

### Reference genome assembly and annotation

Because there was no reference genome available for the *Alyssum montanum-A. repens* complex, we first generated a reference assembly of a diploid *Alyssum gmelinii*. To gain insights into genome structure and complexity, we performed k-mer frequency analysis, estimating the genome size to 608 Mb (consistent with the 622 Mb estimate obtained from flow cytometry), and heterozygosity to 1.4% (Supplementary fig. S3). HiFi reads were assembled using hifiasm (Cheng *et al*., 2021) and the resulting primary assembly (Supplementary table S1) was refined by purging haplotigs, low-quality contigs and organellar DNA. After purging, the quality value (QV) estimated by Merqury tool (Rhie *et al*., 2020) increased from 58.49 to 65.74 (Supplementary table S1, Supplementary fig. S4). The final assembly consists of 168 contigs with total length of 683 Mb, N50 of 12.3 Mb, L50 of 17 and 96.2 % complete BUSCOs scores (Supplementary table S1). The repeat analysis, conducted with RepeatModeler and RepeatMasker, identified 72.09% of the genome as repetitive, including 48.46% of retroelements and 2.6% of DNA transposons. Functional annotation of the reference genome, based on a combination of alignments of protein sequences from the OrthoDB database and newly generated RNA-seq data, resulted in a total of 32,073 annotated protein-coding genes. Using OrthoFinder and BLAST, we assigned *Arabidopsis thaliana* orthologues to 29,266 *Alyssum gmelinii* genes.

### Repeated adaptation to serpentine soil within two ploidies of *Alyssum gmelinii*

Principal component analysis (PCA) of the ionomic composition of the locally collected soil at each population site clearly separated serpentine from non-serpentine soils (along PC1, explaining 44.8% of the variation, Supplementary fig. S5). The main elements contributing to the separation along the PC1 are Co (explaining 20% of the variation), Ni (19%) and Mg (14%), being in higher concentrations in serpentine soil, and Ca (16%), having higher values in non-serpentine soil (Supplementary fig. S5). These results follow the typical expectations for serpentine soil composition.

To test for repeated serpentine colonization in the four sampled geographically proximate population pairs of *Alyssum* (Figure 1A, Supplementary Data S1), we first examined their demographic parameters, relatedness and genetic structure. We sequenced 10 individuals from each population of serpentine (S) and non-serpentine origin (N), obtaining a set of 1,082,883 filtered putatively neutral fourfold-degenerate (4dg) SNPs, with an average sequencing depth of 32x and 4.5% missing data. Site frequency spectra (SFS) exhibit allele frequency patterns consistent with autopolyploidy (Supplementary fig. S6), i.e. lack of over-representation of alleles of intermediate frequency that would otherwise be expected for fixed heterozygotes in allopolyploids (Blischak *et al*., 2023). Allele frequency covariances, calculated using TreeMix (Fig. 1B, Supplementary fig. S7), together with the principal component analysis (PCA) of individual genotypes (Fig. 1C), concordantly split the samples first by species along PC1, and then by region (and ploidy) along PC2. Similarly, Bayesian clustering using Entropy (Shastry *et al*., 2021) (Supplementary fig. S8), with the number of clusters set as two (K = 2), resulted in an *A. gmelinii* cluster separated from the *A. spruneri* one. Increasing the number of clusters to three (K = 3) and four (K = 4) further divided regions (and ploidies) within *A. spruneri* species and within *A. gmelinii*, respectively. Overall, all populations within one geographic pair are genetically fairly closely related (average within-pair Rho = 0.17), with the Austrian pair 2 showing the highest differentiation between S and N populations (Rho = 0.27) (Supplementary fig. S9). Additionally, the pair 2 populations also display the lowest within population nucleotide diversity (ν) (Supplementary Data S1). None of the populations deviate from the expectation of neutrality as indicated by Tajima’s D (range −0.15 – 0.98; Supplementary Data S1). In summary, the populations group by geographic proximity and not by soil type, suggesting repeated colonization of serpentine islands within each species and region without dramatic population size fluctuations.

Serpentine soil presents a challenging environment for plants, often resulting in strong adaptive divergence between colonist and non-colonist populations. By focusing on *A. gmelinii* (pairs 1-2), we aimed to assess potential adaptive differences between populations of serpentine vs. non-serpentine origin when cultivated in their native vs. non-native substrate type through a reciprocal transplant experiment. We observed a significant effect of soil treatment and its interaction with plant substrate of origin on germination rate (generalized linear mixed model) and germination time (mixed-effects Cox proportional hazards model, Fig. 1D) of the seeds. First, seeds generally germinated at a higher rate in non-serpentine treatment (estimate = −1.050, z = −2.735, p = 0.006), with each group germinating at a higher rate in its soil of origin (estimate = 1.675, z = 2.892, p = 0.004). Second, seeds sown in serpentine soil had a significantly higher probability of germinating sooner compared to those in the control soil (HR = 1.398, 95% CI: [1.1, 1.8], p = 0.007). Plant origin alone did not significantly influence the time of germination, however, the interaction between soil of treatment and substrate of origin was highly significant (HR = 2.844, 95% CI: [2, 3.9], p < 0.001, plants of serpentine origin germinated on average about nine days earlier when grown in serpentine soil). Furthermore, plants grown in serpentine soil exhibited significantly smaller rosettes compared to those in the control soil (linear mixed-effects model, β = −2.037, SE = 0.169, t = −12.041, p < 2e-16), however, this was less pronounced for plants of serpentine origin (significant soil treatment by origin interaction effect, β = 1.464, SE = 0.200, t = 7.311, p = 6.03e-12). Final plant biomass was significantly correlated with rosette area (r=0.84, p < 2.2e-16, n=66; Supplementary fig. S10A) and showed a similar pattern of significant soil treatment by origin interaction (β = 1.290, SE = 0.586, t = 2.203, p = 0.0321), overall demonstrating advantage of originally serpentine plants when grown in serpentine substrate (Supplementary fig. S10B).

Finally, we evaluated elemental uptake into roots and leaves of the plants harvested at the end of the experiment. Note that, given the high mortality and very low biomass available of the survived non-serpentine origin plants grown in the S treatment (95% of cultivated N plants died compared to 35% S plants), we lack data regarding this population-treatment combination. In line with the expectation of adaptive response to soil elemental composition, we found that plants of serpentine origin modulated the amount of Ca/Mg ratio uptake relatively to the soil availability (Fig. 1G), increasing it twofold when grown in S soil compared to NS soil treatment. Specifically, originally serpentine plants exposed to serpentine soil drastically increased the uptake of Ca, reaching similar levels of intra-cellular available Ca as observed for the same population in non-serpentine conditions (Supplementary fig. S11-S12B). However, also the uptake of Mg was increased in S soil, therefore decreasing the absolute Ca/Mg ratio value present in the tissues (Supplementary fig. S11-S12). Similarly, Cr was also over-absorbed by plants in serpentine conditions. On the contrary, Ni and Co were maintained at a lower intra-tissue concentration compared to their soil abundance, possibly implying plants’ active exclusion of these elements (Fig. 1H).

In conclusion, assessment of genetic structure suggests repeated colonization of serpentine soil in both investigated *Alyssum* species and regions. In *A. gmelinii*, colonization of serpentine soil was accompanied by substrate adaptation through mechanisms involving differential germination time and ion uptake.

### Genomic signatures of selection at genes relevant for serpentine response

Using the four instances of serpentine colonization in our *Alyssum* dataset, we applied three divergence-based selection scan tools to detect shared footprints of selection associated with repeated serpentine adaptation in the *Alyssum montanum-A. repen*s complex. First, for each population pair we identified the top 1% 1 kb SNP windows of outlier fixation index (F_ST_). This resulted in an average of 1813 outlier windows per population pair that mapped to 338 to 621 candidate genes (depending on the contrast group; Supplementary Data S2). Considering that such an approach does not formally differentiate outliers resulting from directional selection from those caused by other processes such as genetic drift, we refined these gene lists using an additional method, Population Branch Excess, PBE. This method measures the degree to which Population Branch Statistics (PBS) exceeds its expected value, where PBS identifies alleles with extreme frequencies with respect to two outgroup populations (Yassin *et al*., 2016). Doing so, PBE allows us to focus on loci that are under positive selection in the focal population (in our case the serpentine populations) but not in the non-adapted ‘outgroup’ populations (in our case the non-serpentine populations). PBE identified a minimum of 392 to a maximum of 609 candidate genes per population pair (Supplementary Data S3). Overlapping the gene lists resulting from these two methods allowed us to retain a conservative set of on average 188 candidate genes per population pair (Fig. 2A), which we refer to as “population pair-specific genes” from now on. The degree of gene reuse, i.e., repeated emergence as candidates in two serpentine populations, was greater than expected randomly in all combinations (Fisher’s exact test p-value < 0.002, Fig. 2B and Supplementary fig. S13), suggesting some degree of common genomic basis of serpentine adaptation across *Alyssum* ploidies and species. This is consistent with the results of Gene ontology (GO) enrichment analyses of population pair-specific gene lists, which showed a great overlap in biological functions associated with the candidate genes, particularly for biological processes involved in ion transport and homeostasis (Fig. 2D and Supplementary Data S4).

**Fig. 2.**
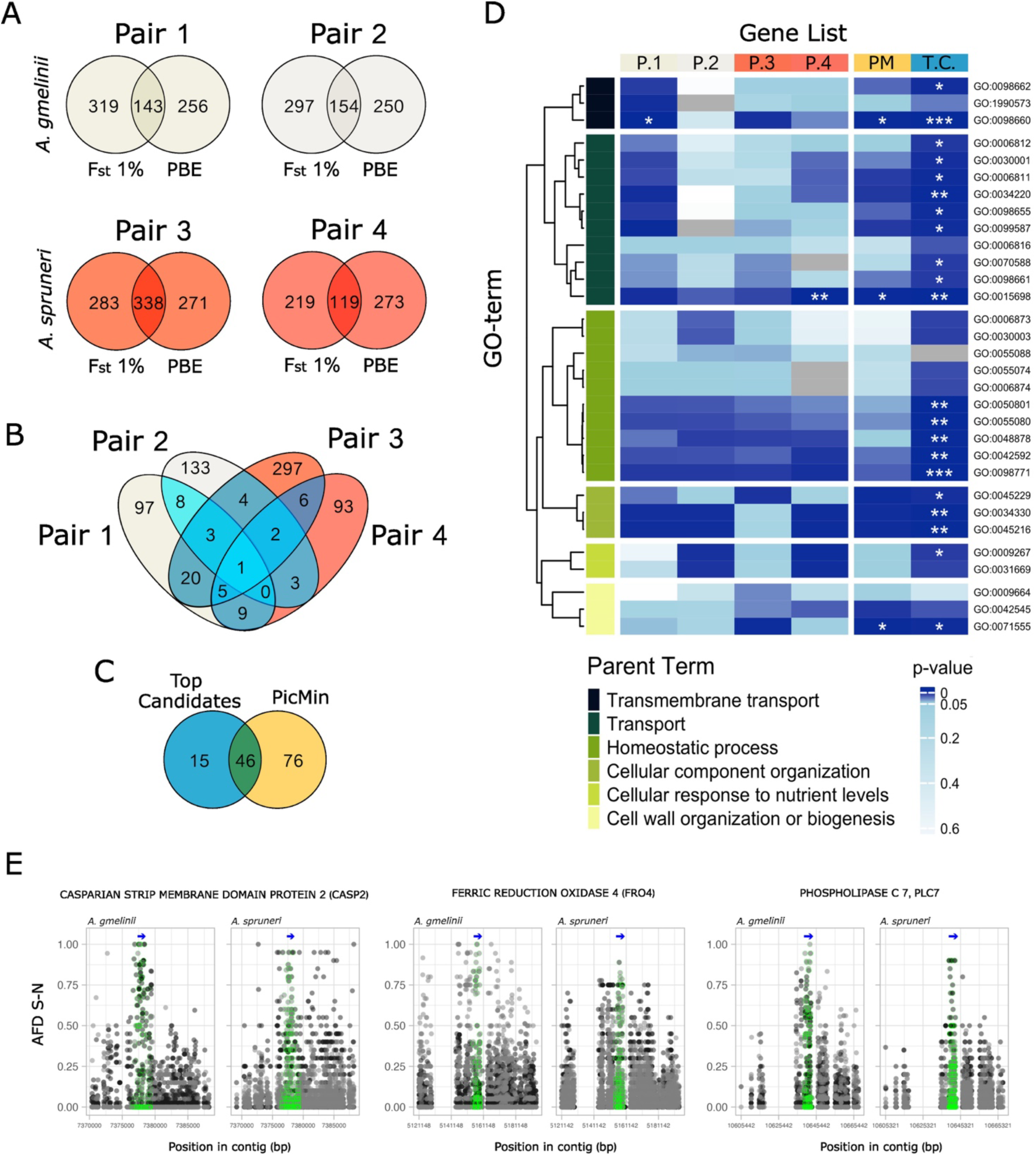
Candidate genes for serpentine adaptation in *Alyssum* and their functions. (A) Venn diagrams showing the number of overlapping genes which were found in the two window-based selection scans (1% outlier F_ST_ and population branch excess, PBE) for each serpentine–non-serpentine population pair. We refer to the overlapping genes as “population pair-specific” candidate genes. (B) Overlap between the population pair-specific candidate adaptive genes resulted in 61 “top candidates” that were repeatedly found as selection candidates in at least two pairs (blue intersections). (C) Number of candidate adaptive genes found both as top candidates and the PicMin results. The overlap is highly significant (Fisher’s exact test p-value < 0.0001). (D) Hierarchically clustered heatmap showing gene ontology (GO)-terms enrichment significance (p-value) for each candidate gene list (population pair-specific – P.1-P.4; PicMin = PM; top candidates = T.C.). Individual GO-terms are clustered and coloured by their parent term using the row annotation. Significance stars refer to the p-values adjusted for multiple testing using the Benjamini-Hochberg correction. (E) Allele frequency difference (AFD) between serpentine (S) and non-serpentine (N) populations for three candidate genes detected as both top candidate and PicMin candidate. Candidate SNP dots falling in the genic region are highlighted in green. A simplified gene model is represented by the blue arrow.

In order to explore genetic parallelisms across *Alyssum* species and to further minimize potential effects of drift and local site-specific selection in each population pair we further focused on a set of genes that would represent a consensus between the two *Alyssum* species in our dataset. Using the gene lists described above, we identified 61 population pair-specific candidate genes that were consistently outliers in at least two population pairs (blue intersections in Fig. 2B, Supplementary Data S5), hereafter referred to as “top candidates”. GO enrichment of *A. thaliana* ortholog genes of such top candidates (58 *A. thaliana* orthologs were found for the 61 candidate genes) well summarizes what was observed for the individual population pairs, identifying significant “biological processes” GO-terms considered important in serpentine adaptation (Arnold *et al*., 2016; Konečná *et al*., 2021), such as inorganic ion homeostasis and transmembrane transport, cell wall organization and regulation of seed growth (Fig. 2D and Supplementary Data S6). Finally, we inferred a complementary list of *Alyssum* consensus candidate genes with the use of PicMin (Booker *et al*., 2023), an order statistics-based method which provides statistical confidence value for each genomic window quantifying the non-randomness of differentiation across multiple lineages (here being the four repeated population pairs in our dataset). In this way, we identified 412 significantly differentiated 1-kbp windows (FDR-corrected q-value < 0.05) which mapped to 122 coding genes (“PicMin” genes - 116 *A. thaliana* orthologs were found for 121 out of 122 candidate genes - Supplementary Data S7). A large proportion of these PicMin genes overlapped with the previously identified top candidates (Fig. 2C), more than what expected randomly (Fisher’s exact test p-value = 1.923087e-102), and shared with them enriched biological functions (Fig. 2D, Supplementary Data S8).

### Evidence for genomic convergence in serpentine adaptation between *Alyssum* and *Arabidopsis arenosa*

Finally, we investigated the potential genetic and functional convergence in serpentine soil adaptation between the *Alyssum montanum-A. repen*s complex and *A. arenosa*, i.e. the only plant species for which a genome-wide list of serpentine adaptive candidates, inferred from individually resequenced genome-wide data, is available (Arnold et al., 2016, Konečná *et al*., 2021). Using the list of 222 *A. thaliana* orthologs identified as parallel serpentine candidates from Konečná *et al*. (2021) (Supplementary Data S9), we tested for gene reuse with our *Alyssum* candidate genes (Fig. 3A). Fisher’s exact tests revealed significant genetic overlap between *A. arenosa* candidate genes and both *Alyssum* top candidates (ten shared genes, p < 0.0001) and the PicMin candidates (11 shared genes, p < 0.0001), supporting the hypothesis of shared targets for positive selection in response to serpentine environments (the thirteen shared candidate adaptive genes are reported with their description in Table 1). Similarly, we observed a functional overlap of such candidate genes, as significantly enriched GO-terms inferred for each species are shared between *Alyssum* and *Arabidopsis* more than what would be expected by chance (*A. arenosa* - *Alyssum* top candidates GO-terms overlap n = 22, p < 0.0001; *A. arenosa* - *Alyssum* PicMin candidates GO-terms overlap n = 15, p < 0.0001; Supplementary fig. S14). To explore in greater detail possible functional convergence, we constructed a protein-protein interaction network (PPIN) using *Arabidopsis thaliana* orthologs in STRING database (Szklarczyk *et al*., 2023), focusing on direct (physical and functional) interactions between *Alyssum* (127, union of top candidates and PicMin candidates) and *A. arenosa* (218) candidate proteins. The resulting network comprised 257 nodes (111 *Alyssum* protein orthologs directly interacting with at least one of 155 *A. arenosa* protein orthologs and vice-versa) and 1,232 edges. A one-sided permutation test confirmed that this level of protein-protein interaction is greater than what would be expected by chance (p = 0.0046, permutation test with 5,000 randomly generated gene lists). We note that the actual connectivity might be even higher, as our approach is focused only on “between-species” direct edges and therefore does not account for “within-species” intermediate nodes, possibly underestimating the true number of “between-species” protein interactions. These results confirm a notable level of adaptive functional overlap between *A. arenosa* and *Alyssum*, reflecting shared molecular networks (Szklarczyk *et al*., 2023) underlying serpentine soil adaptation. GO enrichment analysis of all interacting nodes identified many significant (p-value < 0.05) terms associated with relevant serpentine adaptation processes, that encompassed candidate genes from both groups (Supplementary Data S10). Specifically, we identified a large cluster of “between-species” interacting proteins involved in homeostatic processes, transmembrane transport, transport and cellular responses to nutrient levels (Fig. 3B). Here, genes such as PHT1;1, GORK and NRT2:1 represent possible key hubs. Further relevant clusters were found also for genes related to metabolic processes, cytokinesis, cell cycle, response to stimuli and oxidative stress, suggesting there is a shared importance of such biological processes in mitigating serpentine stress (Fig. 3C-D).

**Fig. 3.**
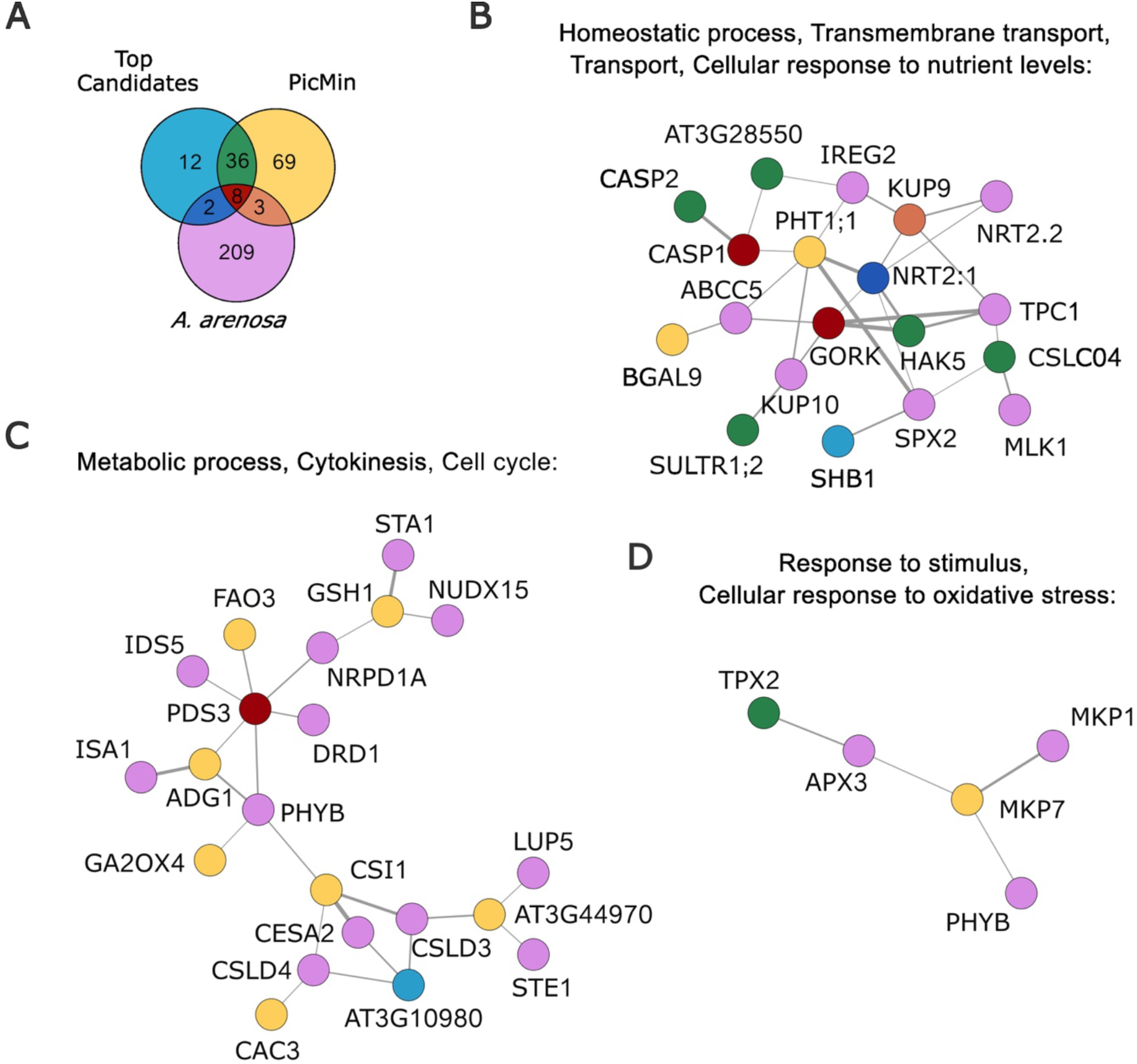
Genetic and functional convergence between *Alyssum* and *Arabidopsis arenosa* serpentine candidate adaptive genes. (A) Venn diagram showing the intersection of *A. thaliana* orthologs for *Alyssum* top candidates, *Alyssum* PicMin candidates, and *A. arenosa* candidate genes (Konečná *et al*., 2021). Shared genes between all three gene lists are highlighted in red. (B–D) Sections of the full functional protein-protein interaction networks (PPIN) constructed from *A. thaliana* orthologs of candidate genes, focusing on between-species direct interactions. Nodes represent proteins belonging to *Alyssum* or *A. arenosa*, or both, based on colour codes as in A. Each node is labelled with its corresponding gene name. Edges represent protein interaction, and their thickness reflects the interaction combined score (i.e. the interaction confidence). (B) Interactions between nodes related to processes such as homeostasis, transmembrane transport, transport, and cellular responses to nutrient levels. (C) Interactions between candidate genes involved in metabolic processes, cytokinesis, and the cell cycle. (D) Network showing genes associated with responses to stimuli and oxidative stress.

**Table 1.** List of the thirteen genes shared between *Alyssum* and *Arabidopsis arenosa* as candidate genes to serpentine adaptation, with their functional description.

| *A. gmelinii* ID | *A. lyrata* ID | *A. thaliana* ID | Description |
| --- | --- | --- | --- |
| Ag007G0893600 | AL7G42140 | AT4G14220 | RING-H2 GROUP F1A (RHF1A) |
| Ag007G0893500 | AL7G42150 | AT4G14210 | PHYTOENE DESATURASE 3 (PDS3) |
| Ag007G0893900 | AL7G42110 | AT4G14230 | MAGNESIUM RELEASE 2, MGR2 |
| Ag004G0610800 | AL8G37350 | AT5G60760 | P-loop containing nucleoside triphosphate hydrolases<br>superfamily protein |
| Ag007G0893700 | AL7G42130 | AT4G14225 | A20/AN1-like zinc finger family protein |
| Ag007G0896500 | AL7G42240 | AT4G14147 | ARPC4 |
| Ag034G3751600 | AL1G18450 | AT1G08090 | NITRATE TRANSPORTER 2:1 (NRT2:1) |
| Ag054G4879000,<br>Ag054G4878900 | AL4G32190 | AT2G36100 | CASPARIAN STRIP MEMBRANE DOMAIN<br>PROTEIN 1 (CASP1) |
| Ag031G3332900 | AL5G37220 | AT3G55940 | PHOSPHOLIPASE C 7, PLC7 |
| Ag034G3742000 | AL7G46260 | AT5G37500 | GATED OUTWARDLY-RECTIFYING K<sup>+</sup><br>CHANNEL (GORK) |
| Ag012G1282400 | AL2G41520 | AT3G42170 | DAYSLEEPER |
| Ag007G0896300 | AL7G42010,<br>AL7G42020 | AT4G14280 | ARM repeat superfamily protein |
| Ag011G1188000 | AL7G33950 | AT4G19960 | K<sup>+</sup> UPTAKE PERMEASE 9 (KUP9) |

## Discussion

Here we leveraged four serpentine colonizations in two closely related *Alyssum* species (*A. gmelinii* and *A. spruneri*) and identified adaptive candidate genes responsible for adaptation to these extreme soils. We further investigated the extent of evolutionary repeatability not only between the two *Alyssum* species, but also between members of two genera representing distinct tribes of the Brassicaceae family (*Alyssum*, Alysseae and *Arabidopsis*, Camelineae). Our results provide evidence of both genetic and functional convergence in the genomic basis of serpentine adaptation between those genera, shedding light on the mechanisms driving adaptive convergence in extreme environments.

### Repeated serpentine colonization and adaptation

A necessary prerequisite of an efficient genome scan for selective sweeps is a deeper knowledge of the underlying population genetic structure. Our analyses suggest that serpentine *Alyssum* populations represent independent colonization events within each sampled species and region. This pattern aligns with findings in other systems, where parallel colonization of serpentine sites from surrounding non-serpentine areas seems to be the rule in the majority of species that occupy both substrate types (e.g. *Odontarrhena bertolonii* (Mengoni *et al*., 2003), *Arabidopsis arenosa* (Konečná *et al*., 2021), *Cerastium alpinum* (Berglund and Westerbergh, 2001), *Galium pusillum* agg (Knotek and Kolář, 2018), *Lasthenia californica* complex (Rajakaruna *et al*., 2003), *Silene paradoxa* (Lazzaro *et al*., 2021), *Solidago virgaurea* (Sakaguchi *et al*., 2019)). Despite repeated independent colonization, populations of both substrate types exhibit comparable genetic diversity and they are devoid of signals of strong population bottlenecks. Such observation suggests serpentine colonization was not accompanied by drastic founder events and it rather aligns with a scenario of consistent gene flow maintaining low inter-population differentiation genome-wide (Rho). Similar situations have been observed also in *Odontarrhena serpyllifolia* (Sobczyk *et al*., 2017), *Arabidopsis arenosa* (Arnold *et al*., 2016; Konečná *et al*., 2021), tetraploids of *Biscutella laevigata* (Bürki *et al*., 2024), *Helianthus exilis* (Sambatti and Rice, 2006), *Knautia arvensis* (Kolář et al., 2012) and *Silene paradoxa* (Lazzaro *et al*., 2021). Nevertheless, our study adds to the body of evidence showing that even in the presence of gene flow, selection can maintain differentiation at traits critical for survival and reproduction (Gonzalo-Turpin and Hazard, 2009; Fitzpatrick *et al*., 2015; Arnold *et al*., 2016; Tigano and Friesen, 2016; Wang *et al*., 2020; Zhang *et al*., 2021).

We detected a strong signal of serpentine adaptation in a transplant experiment involving *A. gmelinii*. Firstly, our results revealed that seeds sown on serpentine soil germinated earlier than those sown on non-serpentine soil, with populations of serpentine origin exhibiting the earliest germination overall. These patterns suggest germination timing might be an important adaptive trait for survival and fitness in serpentine habitats, as similar results have been detected also in *Arabidopsis lyrata* (Veatch-Blohm *et al*., 2017). We speculate that early germination on serpentine soil could provide a critical advantage by allowing seedlings to establish before the onset of harsh conditions, such as drought or nutrient limitations, which are characteristic of serpentine environments (Brady *et al*., 2005; Harrison and Rajakaruna, 2011). Such an advantage of early germination timing has been reported to be a key mechanism of local adaptation in other systems, including *Arabidopsis thaliana* (Postma *et al*., 2016; Postma and Ågren, 2022) and *Streptanthus tortuosus* (Gremer *et al*., 2020). We also detected a substrate-of-origin specific effect of soil treatment for rosette area and biomass, which could be partially explained by the observed differential elemental accumulation in plant tissues. Similarly to what was described for serpentine *Alyssum* (Tomović *et al*., 2013), *Odontarrhena lesbiaca P. Candargy* (Kazakou *et al*., 2010)*, Helianthus bolanderi* (Walker *et al*., 1955) and *Arabidopsis arenosa* (Arnold *et al*., 2016), we observed that our serpentine-tolerant populations are able to uptake higher quantities of Ca, despite the depleted levels in the serpentine soil, without accumulating excessive quantities of Mg and therefore exhibit an elevated Ca/Mg ratio in their tissues. Concomitantly, serpentine-adapted plants responded to the elevated levels of Ni in serpentine soil by restricting its uptake to the shoots, as typically observed in non-hyperaccumulator plants (Doubková *et al*., 2012; Palm and Van Volkenburgh, 2014; Kolář *et al*., 2014; Chathuranga *et al*., 2015; Arnold *et al*., 2016; Salehi Eskandari *et al*., 2017; Konečná *et al*., 2021). Absence of hyperaccumulation phenotype in this group is also in line with previous observations in serpentine *Alyssum spruneri* from Serbia (Tomović *et al*., 2013) and non-serpentine *A. gmelinii* coastal populations (Ievinsh *et al*., 2020). However, it should be noted that despite lower than in the cultivation soil, the average level of Ni tissue accumulation in our S-treated plants was still high, 82 mg kg^-1^ on average, with many individuals showing an accumulation level > 100 mg kg^-1^. This is a notable level of Ni concentration, enough to classify our *A. gmelinii* plants as strong accumulators (Brooks *et al*., 1979), similarly as it was observed for *Alyssum spruneri* from Serbia (Tomović *et al*., 2013). Interestingly, also Cr, despite being toxic for plants, was considerably accumulated in *A. gmelinii* tissues, with an average accumulation of 41 mg kg^-1^. These findings indicate that *A. gmelinii* genetically adapted to the challenging serpentine soil composition by a suite of traits that include accumulation and restricted uptake of different elements.

### Genetic basis of serpentine adaptation in *Alyssum*

Investigation of the genetic basis of serpentine adaptation uncovered signals of positive selection on genes responsible for ion transport and homeostasis. For example, the two population pairs of *A. gmelinii* exhibited signatures of selection on genes such as the root expressed high-affinity sulphate transporter 1;1 (SULTR1;1), implicated as candidate locus in previous studies on *Arabidopsis* soil adaptation (Arnold *et al*., 2016; Guggisberg *et al*., 2018; Bontpart *et al*., 2024). Notably, a significant proportion of genes showing signature of positive selection are shared between *A. gmelinii* and *A. spruneri*, suggesting that both species might have gone through a parallel process of local adaptation, sourcing from a common pool of standing variation shared among the species. That such allele sharing is likely in *Alyssum* corresponds with close relationships between both species (they belong to the same species complex) and with recent findings on the complex reticulate phylogeny of the entire group involving repeated whole genome duplications, area shifts and hybridisation events (Španiel *et al*., 2011, 2012, 2023; Melichárková *et al*., 2017; Španiel, Zozomová-Lihová, *et al*., 2017; Španiel, Marhold, *et al*., 2017).

Multiple candidate genes that were shared between both *Alyssum* species, are directly involved in the “Serpentine Syndrome”. Firstly, they encode proteins involved in ion transport, such as SULTR1;1 again, but also SULTR1;2, the high-affinity nitrate transporters NRT2:1 and NRT2.3, the potassium transporters KUP9 and HAK5, the transmembrane magnesium transporter MGT7, the ferric reduction oxidase FRO4, the phosphate transporters PHT1;1 and PHT1;2 and the guard cell outward potassium channel GORK. These genes have been described to be involved in serpentine, siliceous and calcareous soil adaptation in *A. lyrata*, *A. arenosa* and *Microthlaspi erraticum* (Turner *et al*., 2010; Arnold *et al*., 2016; Guggisberg *et al*., 2018; Mishra *et al*., 2020; Konečná *et al*., 2021), metalliferous soil adaptation in *A. halleri* and *A. arenosa* (Preite *et al*., 2019), salinity adaptation in *Brassica fruticolosa* (Busoms *et al*., 2024) and aluminium tolerance in *Brachiaria* grasses and *A. thaliana* (Sawaki *et al*., 2009; Worthington *et al*., 2021). Secondly, they encode proteins related to uptake of nutrients and water, such as CASP1, CASP2 and ESB1, which together are responsible for root uptake selectivity, guiding the Casparian strip formation and therefore endodermal differentiation (Roppolo *et al*., 2011; Kamiya *et al*., 2015). Finally, we observed differentiated genes related to life-history traits, such as SHB1, which control endosperm development and seed size (Zhou *et al*., 2009), EDA4, responsible for embryo development (Pagnussat *et al*., 2005), PUB4, RHF1A, DAYSLEEPER and CSI1, affecting development and fertility (Hauser *et al*., 1995; Liu *et al*., 2008; Wang *et al*., 2013; Knip *et al*., 2013).

### Convergence in genetic basis of serpentine adaptation between Alyssum and Arabidopsis

Given the multiple hazards posed by serpentine conditions, such as lack of nutrients, toxicity and propensity to drought, understanding the evolutionary processes that helped plants to colonize and thrive in such an environment present an opportunity for agronomical planning and conservation efforts. The study of convergent adaptation, especially between fairly divergent species, uncovers the genetic bases of repeated adaptation and therefore enhances our ability to predict new, and old, evolutionary outcomes (Bohutínská and Peichel, 2024). Similarly to our case in two *Alyssum* species, repeated adaptation to serpentine soil within *A. lyrata* and *A. arenosa* involved a conspicuous number of shared genes (Arnold *et al*., 2016; Konečná *et al*., 2021), what is expected for closely related species (Bohutínská and Peichel, 2024). On top of that, however, we also detected a significant genetic reuse between two *Alyssum* species and *A. arenosa*, despite their higher evolutionary divergence of approximately 17-20 Mya (Hendriks *et al*., 2023), exposing the pivotal role of the convergent genes in enabling plants to overcome serpentine challenges. These findings align with studies on convergent evolution in other Brassicaceae systems, such as for alpine adaptation (Rellstab *et al*., 2020), but are in contrast with studies on arctic stresses adaptation (Birkeland *et al*., 2020) and partly also with studies on adaptation to whole-genome duplication (Bohutínská, Alston, *et al*., 2021; Bray *et al*., 2024) which found no or very limited gene-level convergence across Brassicaceae.

Going beyond convergent candidate genes, we also found significant convergence at the level of functional networks by reconstructing patterns of protein-protein interactions between all candidate adaptive proteins across the genera. We found that even non-convergent candidate genes are often directly connected in clusters related to metal homeostasis, transmembrane transport, and cellular stress responses. These findings suggest there might be conserved genetic pathways, at least within the Brassicaceae species, underlying plant adaptation to extreme soil conditions, similarly as it was documented for arctic adaptation (Birkeland *et al*., 2020), alpine (Bohutínská, Vlček, *et al*., 2021) and post whole-genome duplication (Bohutínská, Alston, *et al*., 2021; Bray *et al*., 2024) adaptation in Brassicaceae. Further research, including functional validation of candidate genes and detailed characterization of the adaptive pathways, will be crucial to unravel the precise shared molecular mechanisms driving serpentine adaptation (Konečná *et al*., 2020). Additionally, expanding this approach to other serpentine-adapted species may provide a broader understanding of the generality of these adaptive strategies across the plant kingdom and uncover mechanisms driving evolutionary repeatability in adaptation towards strong environmental challenges.

## Supplementary data

Supplementary Figures and Tables: contains additional result figures and tables and their relative description. Supplementary Data: dataset information, candidate gene lists and their functional enrichment. Supplementary Methods: plants and soil ionomic analysis methods description.

## Acknowledgements

We thank Miroslav Poláček for assistance during fieldwork and Karolína Havlíková and Veronika Vlčková for help performing the transplant experiment and phenotyping.

## Data availability

Sequence data that support the findings of this study are deposited in the NCBI (https://www.ncbi.nlm.nih.gov/bioproject/) under BioProjects PRJNA1226118 (short read data) and PRJNA1220107 (long read data).

