## Supplementary fig. ;Supplementary table S1 for "Genomic basis of adaptation to serpentine soil in two *Alyssum* species shows convergence with *Arabidopsis* across 20 million years of divergence"

**Supplementary figures and tables:**

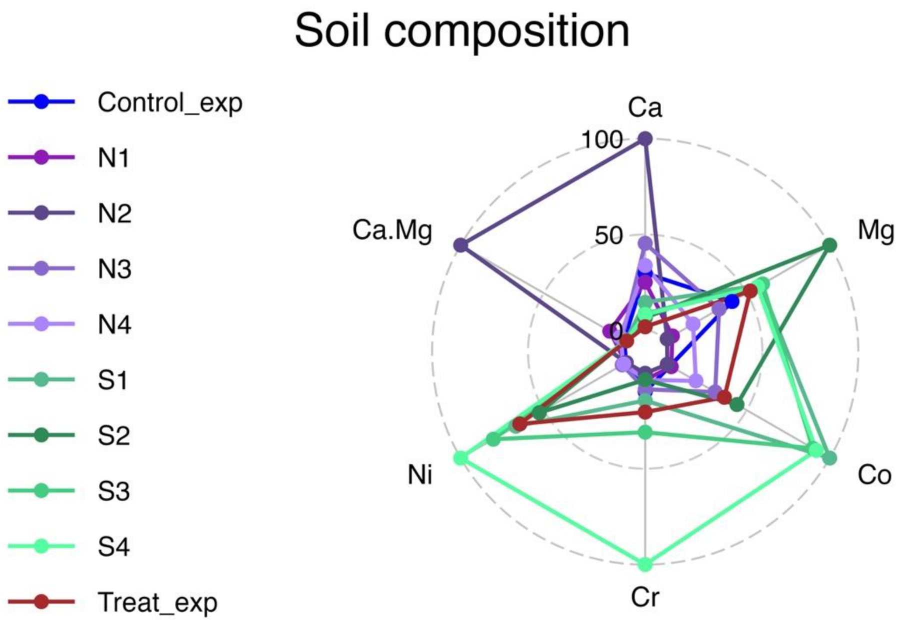


**Supplementary figure S1.** Radar chart of soils elemental composition (mg/kg) in percentage, including calcium (Ca), magnesium (Mg), cobalt (Co), chromium (Cr), nickel (Ni) and ratio of calcium/magnesium (Ca.Mg). Each line represents one population and it is coloured accordingly. Here, we included data about local soil for each population, plus data about the soil used for the transplant experiment – non-serpentine soil (Control_exp) and serpentine soil (Treat_exp). Experimental soils follow similar patterns of soil composition as the respective natural original soils.

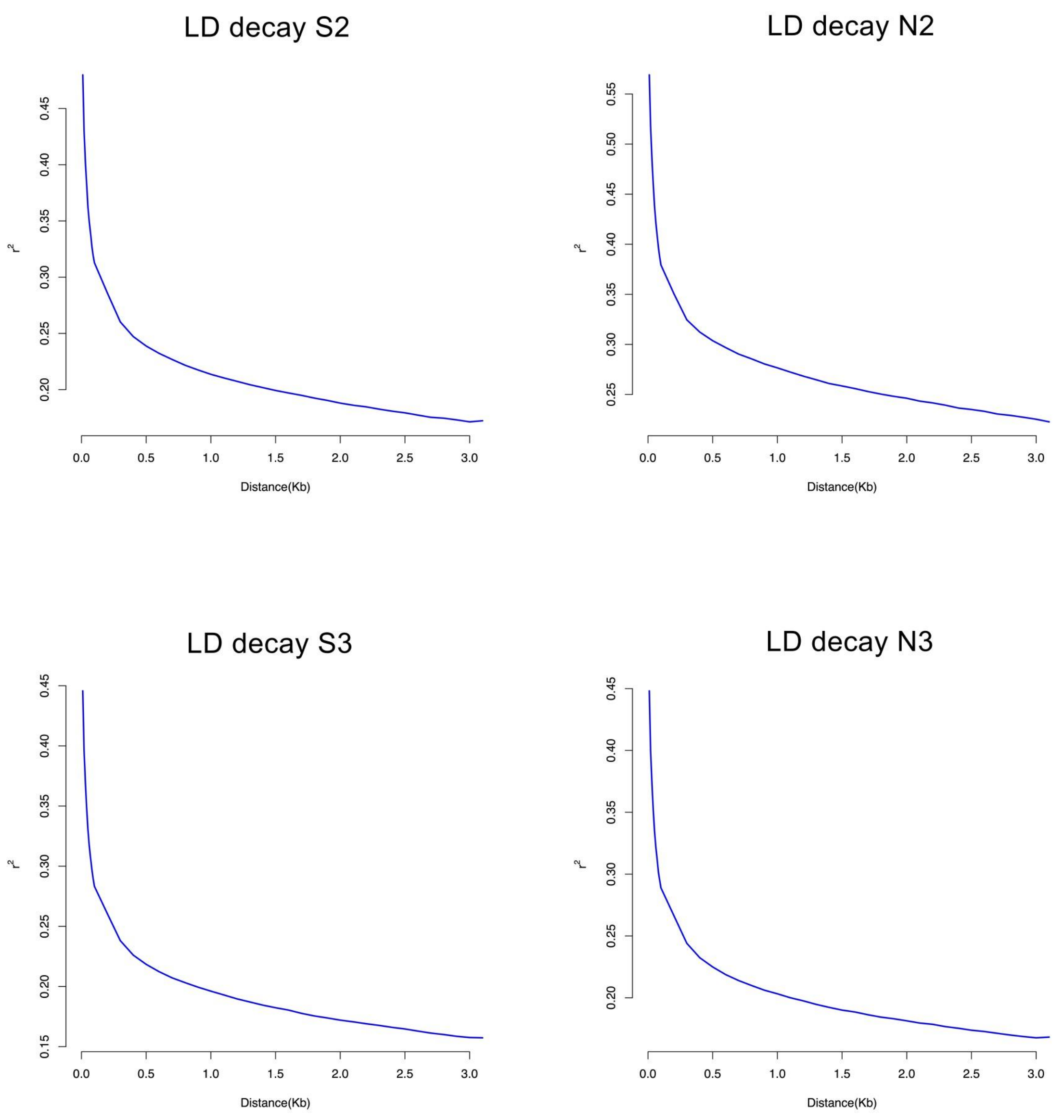


**Supplementary figure S2.** Decay in genotypic correlation ($r^2$), i.e. the proxy for linkage-disequilibrium (LD) calculated in diploid populations. $r^2$ estimates are plotted against the site distance in the reference genome (Kb).

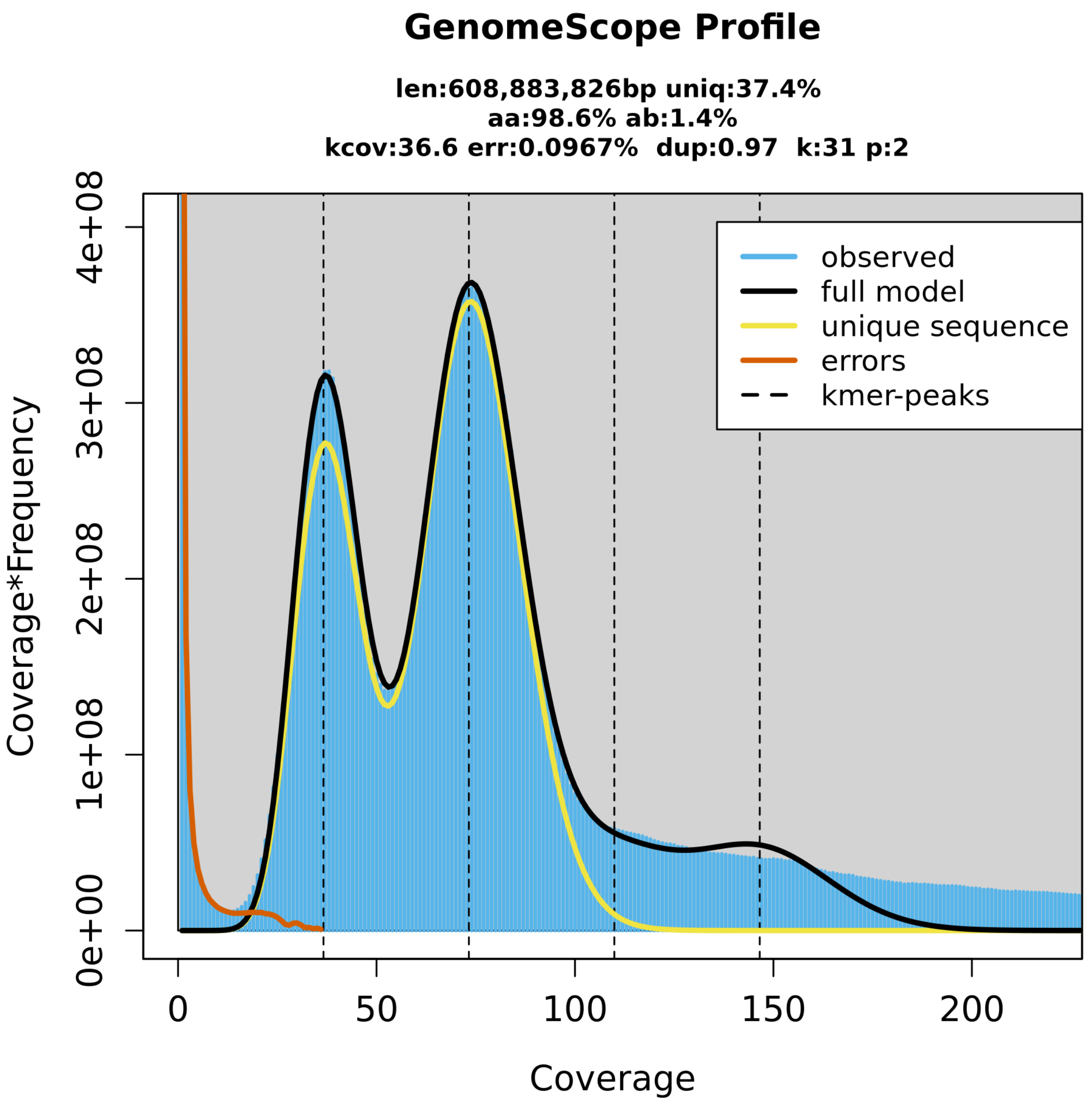


**Supplementary figure S3.** GenomeScope k-mer spectra, based on PacBio HiFi reads, of the *Alyssum gmelinii* genome assembly. Peaks correspond to distinct k-mer coverage levels, indicating haploid and diploid k-mer distributions. Estimated genome size: 608.9 Mbp, with 37.4% unique sequence, 0.97% duplication, and an estimated heterozygosity rate of 1.4%.

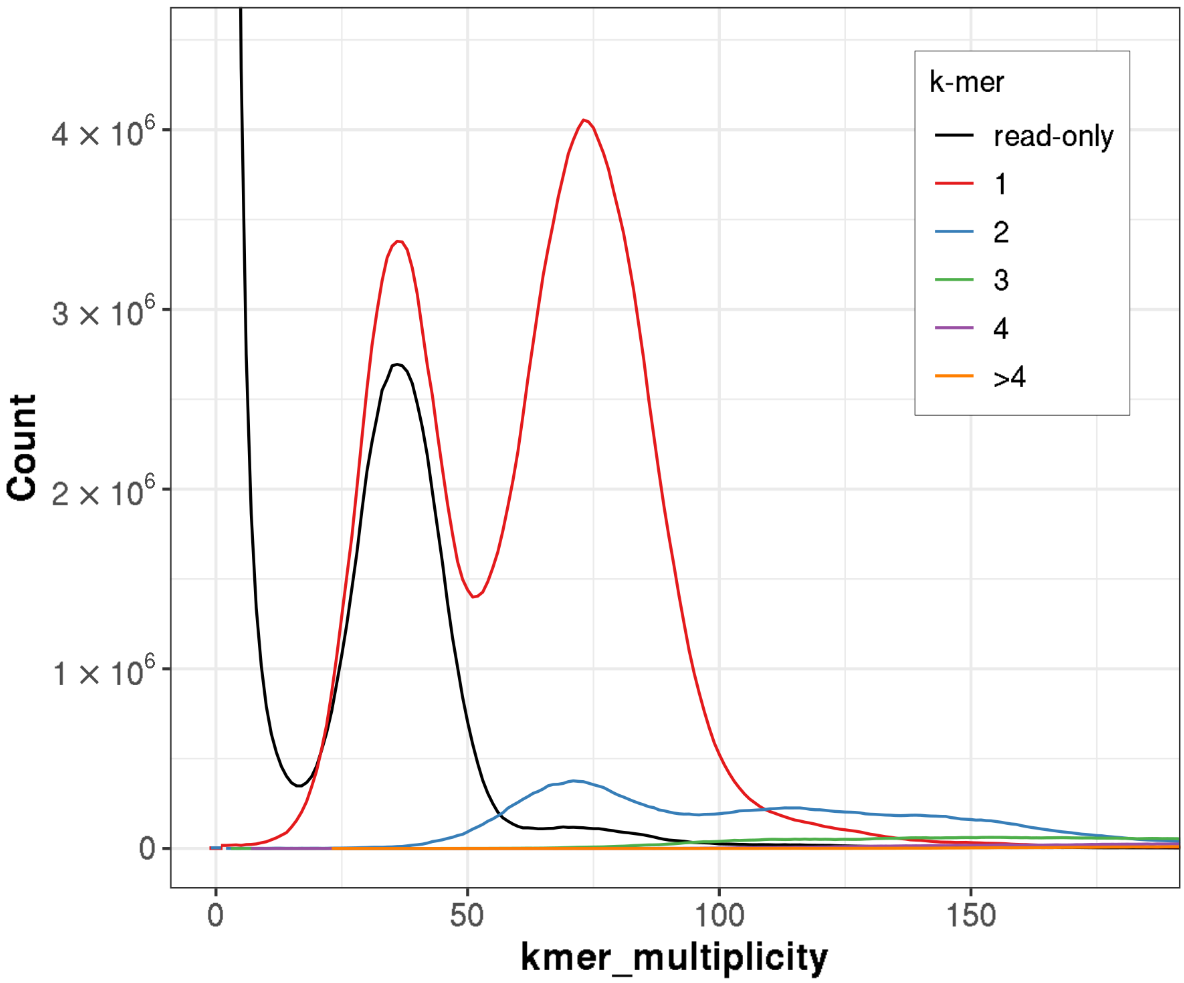


**Supplementary figure S4.** Copy number spectrum (spectra-cn) of the primary assembly of the *Alyssum gmelinii* genome generated using Mercury. The plot illustrates the distribution of *k*-mers across varying multiplicities. The black line represents the *k*-mer frequency derived from sequencing reads, while the colored lines correspond to *k*-mers assigned to distinct copy number categories.

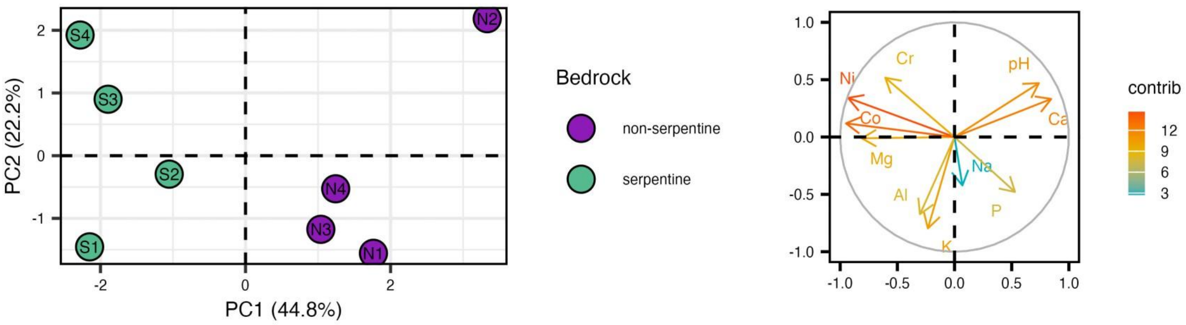


**Supplementary figure S5.** PCA of soil elemental composition for each sampled population on the left. Dots are coloured by soil type and labelled by the population pair name. Variables correlation plot on the right, with loadings of soil variables coloured by their contribution to the PC1, expressed in percentage.

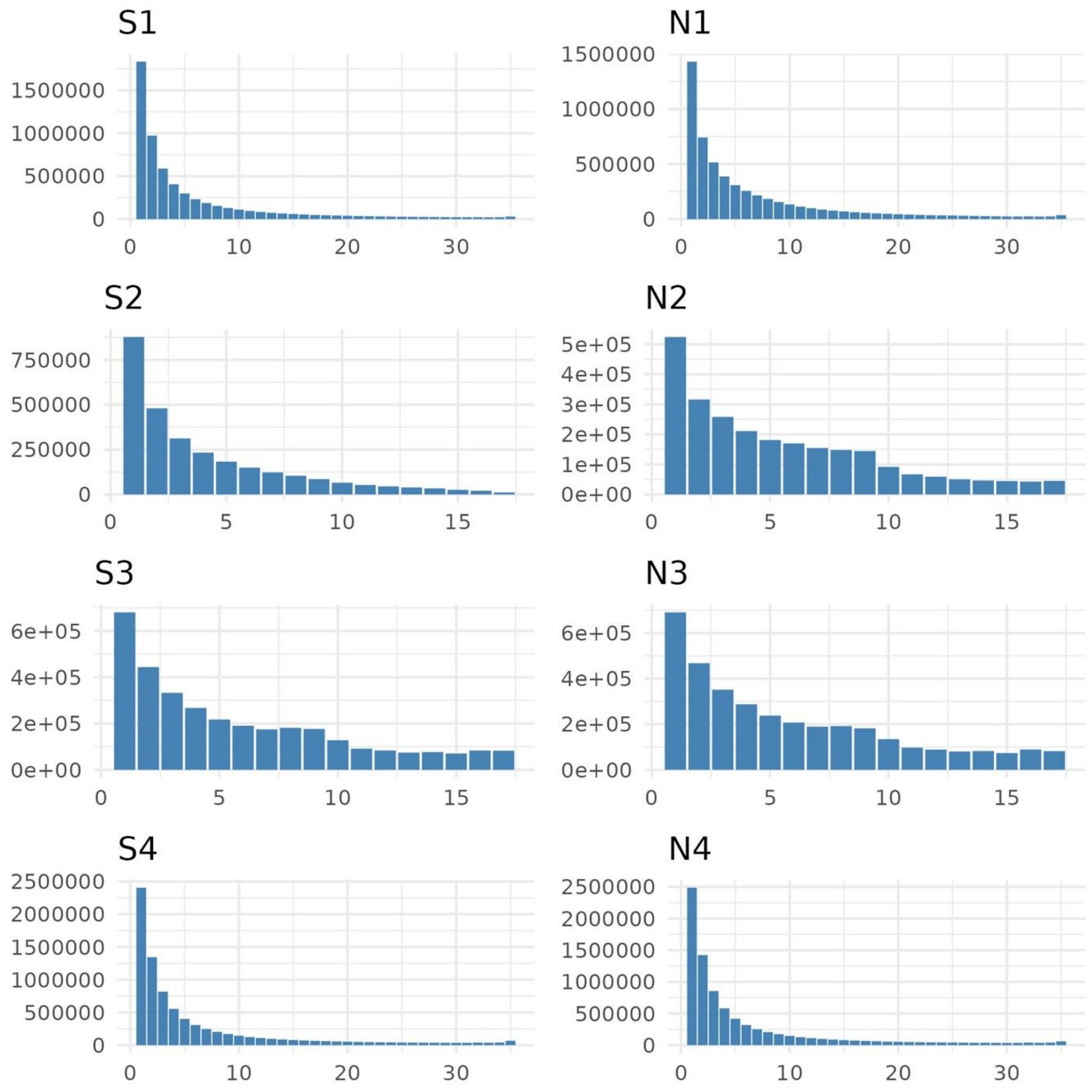


**Supplementary figure S6.** Unfolded site frequency spectrum for individual populations based on 9 individuals (18 and 36 alleles in diploid and autotetraploid populations, respectively). Bars represent the number of alleles found in each allele frequency class. The counts for invariant sites are removed for the sake of visualization.

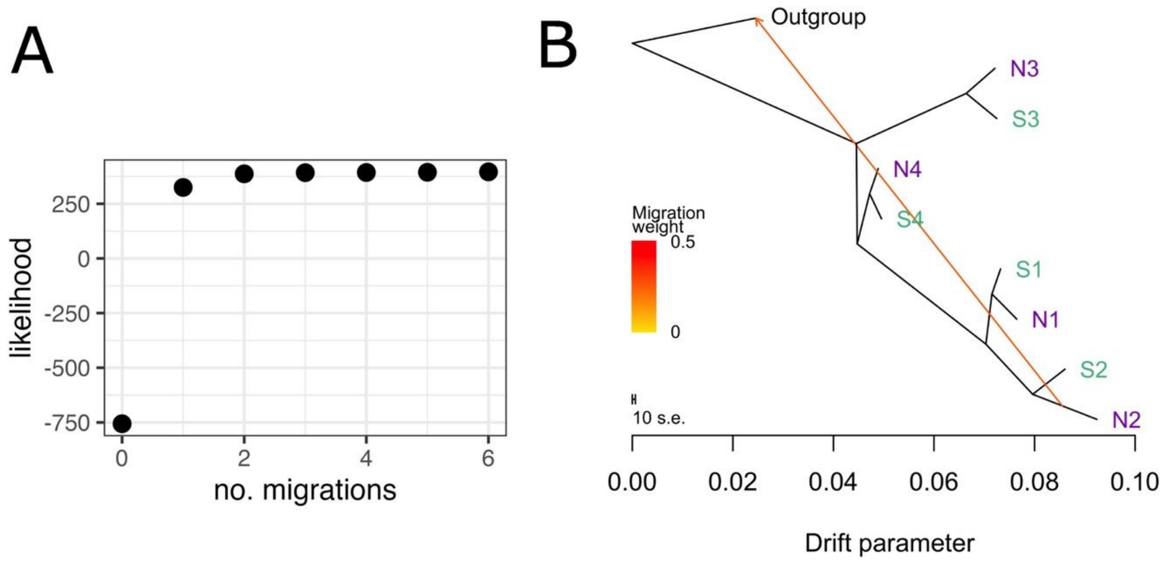


**Supplementary figure S7.** (A) Pattern of increasing tree/network topology likelihood with rising number of migration events in Treemix analyses. A higher than one number of migration edges does not improve the model. (B) Allele frequency covariance graph of the studied populations calculated using neutral fourfold-degenerate SNPs (4dg), including the first migration edge between populations shown with an arrow directing towards the recipient group, coloured according to its weight (ancestry percentage received from the donor). The colour of population labels refers to the soil type of origin (green – serpentine, violet – non-serpentine). The outgroup represents a diploid *Alyssum repens* population from Kleinstübing, Austria.

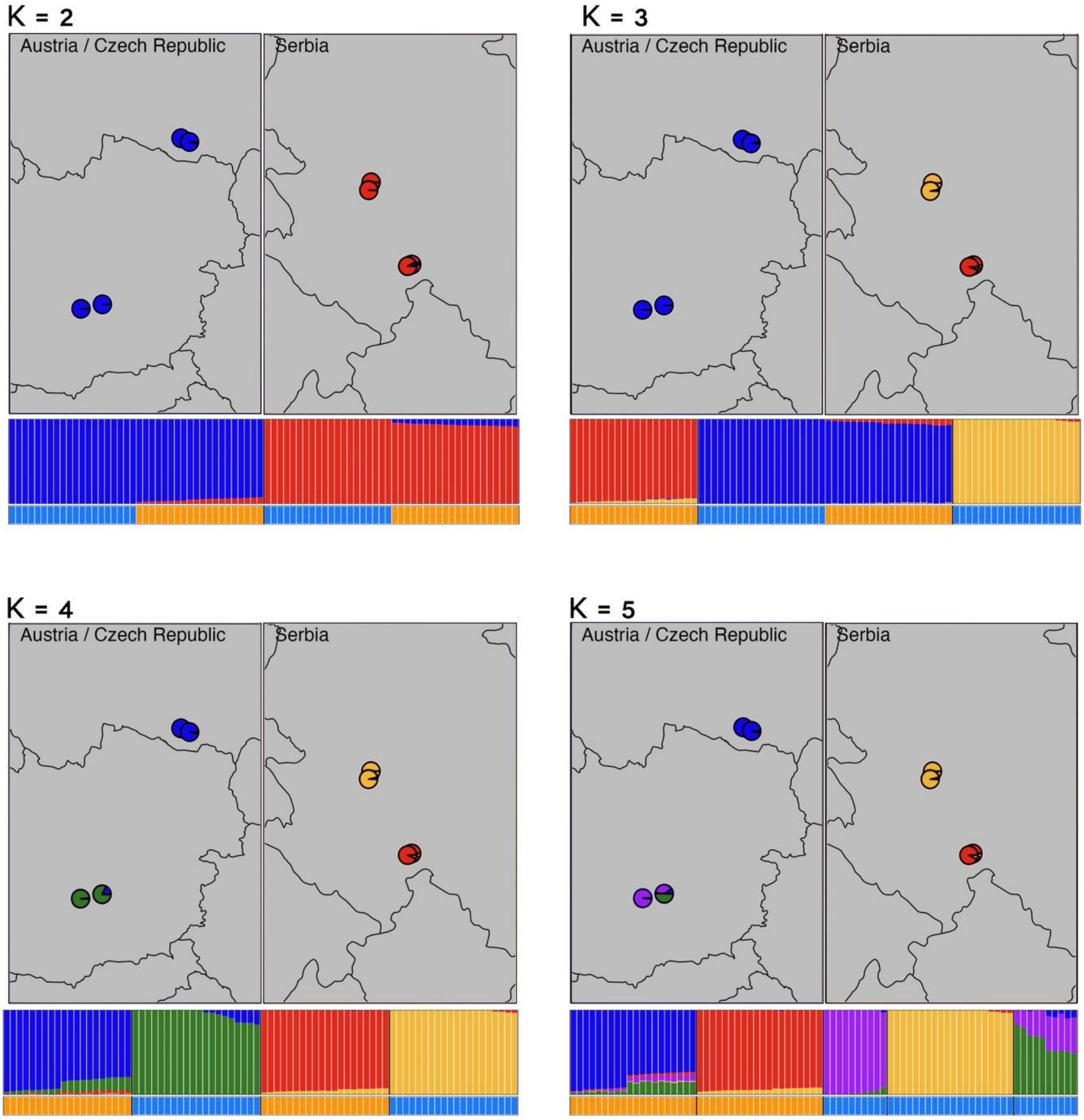


**Supplementary figure S8.** Genetic differentiation of our study system inferred by means of Bayesian population clustering. In each picture we show the geographical distribution of the populations indicated as pie charts reflecting the proportional assignment to particular genetic clusters identified by Entropy, bar plots reflecting individual assignment into each genetic cluster identified by Entropy and the ploidy of each individual (lightblue – diploid, orange – tetraploid). The results are shown for a defined number of clusters (K) ranging from two to five. In line with Treemix analysis, Entropy results split firstly the populations by species (K=2), and secondly by region (and ploidy, K=4).

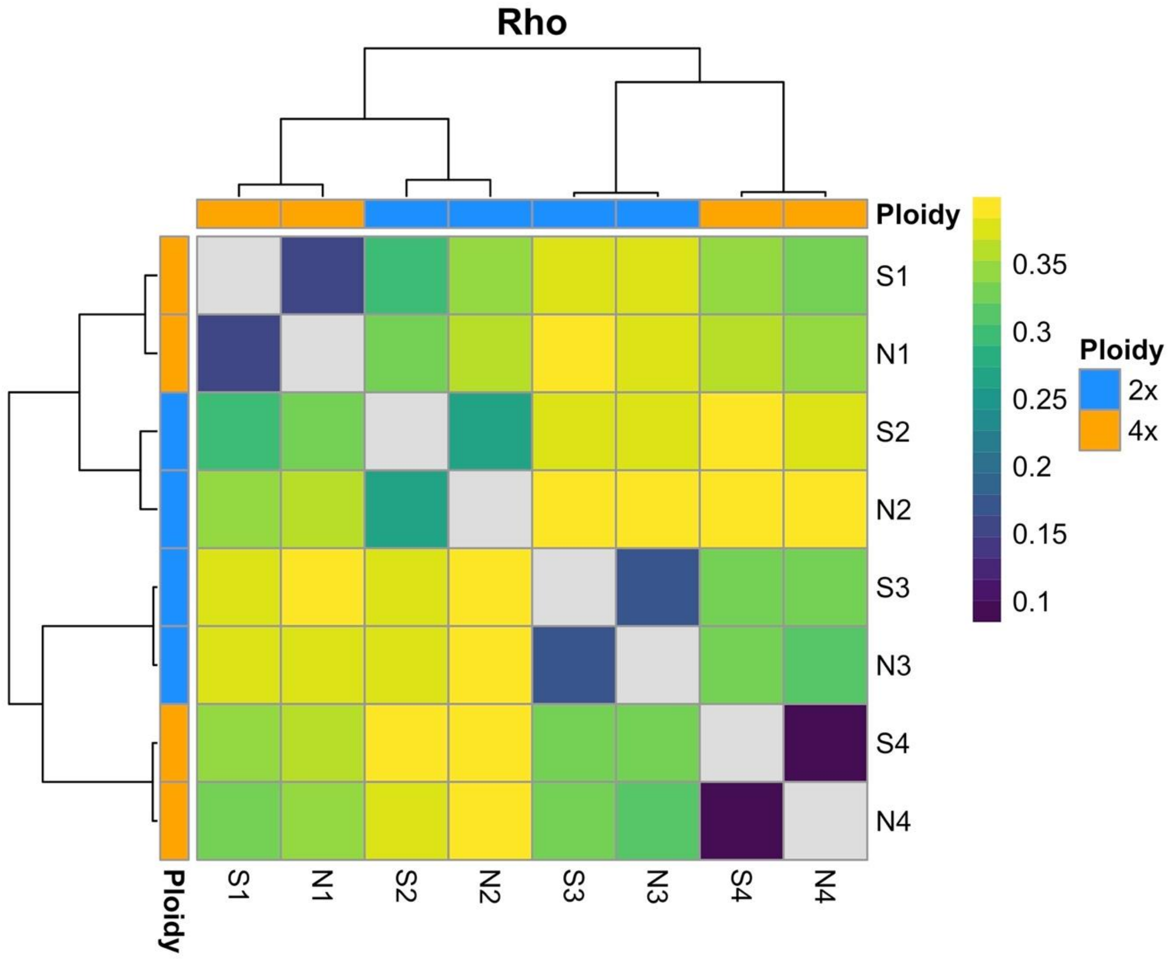


**Supplementary figure S9.** Row and columns hierarchically clustered (following *Ward*'s minimum variance method) heatmap showing the magnitude of genome-wide pairwise genetic differentiation (Rho) between our study populations. High genetic differentiation (Rho) is represented by shadows of yellow, while low genetic differentiation is coloured in blue. Columns and rows are annotated with population name and ploidy.

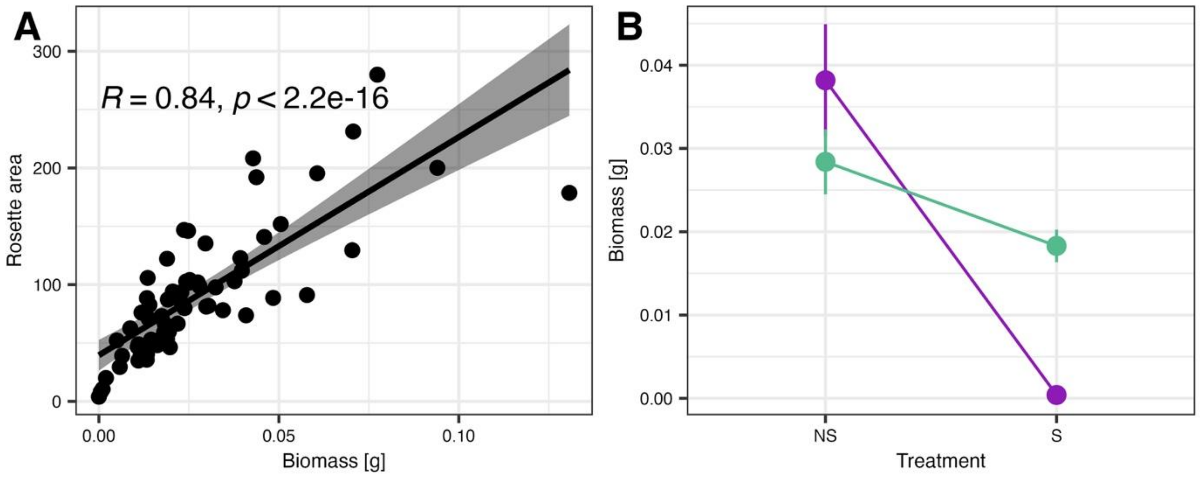

**Supplementary figure S10.** (A) Scatter plot and Spearman rank positive correlation of plants dry biomass (g) at day 67 and rosette area scored at day 25 after repotting. The solid line represents the fitted regression line and the shaded area its standard deviation (r = 0.84, p < 2.2e-16). (B) Differences in the final dry biomass (g) of experimental plants grown in serpentine (S) and non-serpentine (NS) soil conditions. Colours refer to population soil of origin (green – serpentine, violet – non-serpentine). Points denote mean, error bars depict standard error of mean. Linear mixed effect model shows significance of the soil treatment interaction with soil of origin: β = 1.290, SE = 0.586, t = 2.203, p = 0.0321.

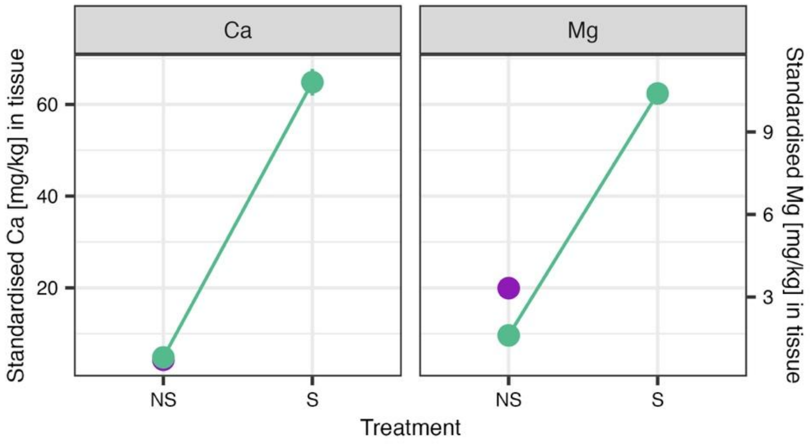


**Supplementary figure S11.** Differences in calcium (Ca) and magnesium (Mg) (mg/kg) uptake in tissues of plants cultivated in S and NS soils (n = 78 individual samples), standardised by the amount of the elements present in the soil treatment. Colours refer to population soil of origin (green – serpentine, violet – non-serpentine). Points denote mean, error bars depict standard error of mean.

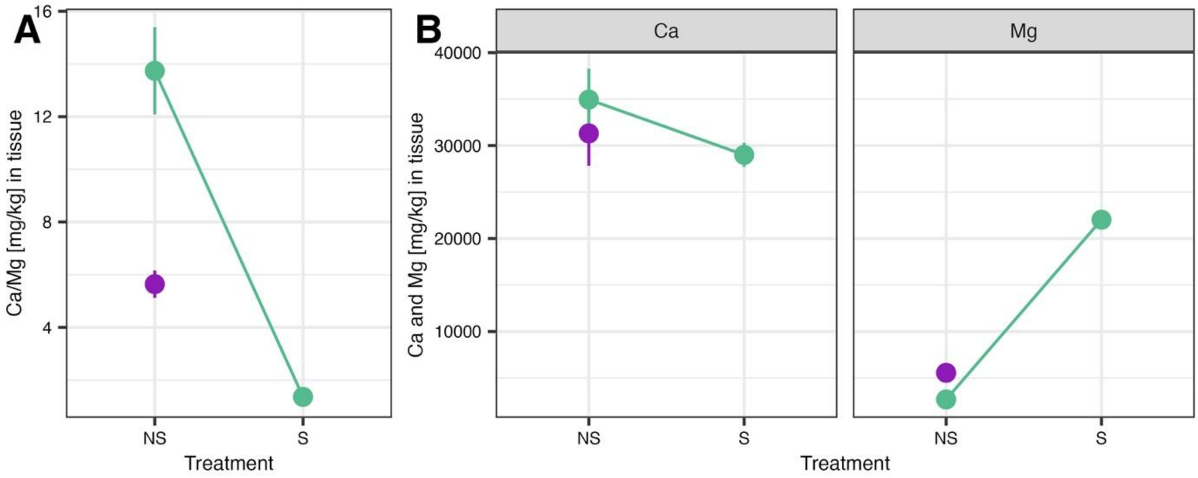


**Supplementary figure S12.** Differences in the absolute uptake of calcium/magnesium ratio (Ca/Mg) (A) and calcium (Ca) and magnesium (Mg) singularly (B) – in mg/kg – in tissues of plants cultivated in S and NS soils (n = 78 individual samples). Colours refer to population soil of origin (green – serpentine, violet – non-serpentine). Points denote mean, error bars depict standard error of mean.

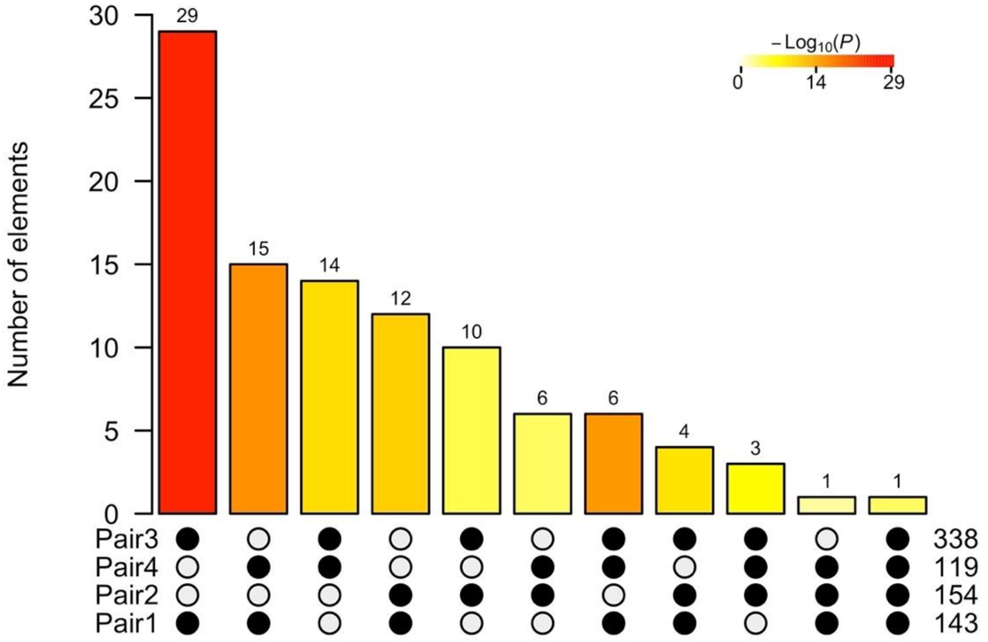


**Supplementary figure S13.** Intersection of population pair-specific gene candidates demonstrating that the number of genes repeatedly found as candidates across two, three, and four population pairs are greater than expected by chance alone (all intersections are significant at $p < 0.002$, one-sided Fisher's exact test). The colour intensity of the bars represents the p-value significance of the intersections, and the number of genes shared is reported on each intersection bar. Notably, the highest numbers of overlapping genes are found between the tetraploid population of *A. gmelinii* (pair 1) and the two populations of *A. spruneri* (pair 3 and 4).

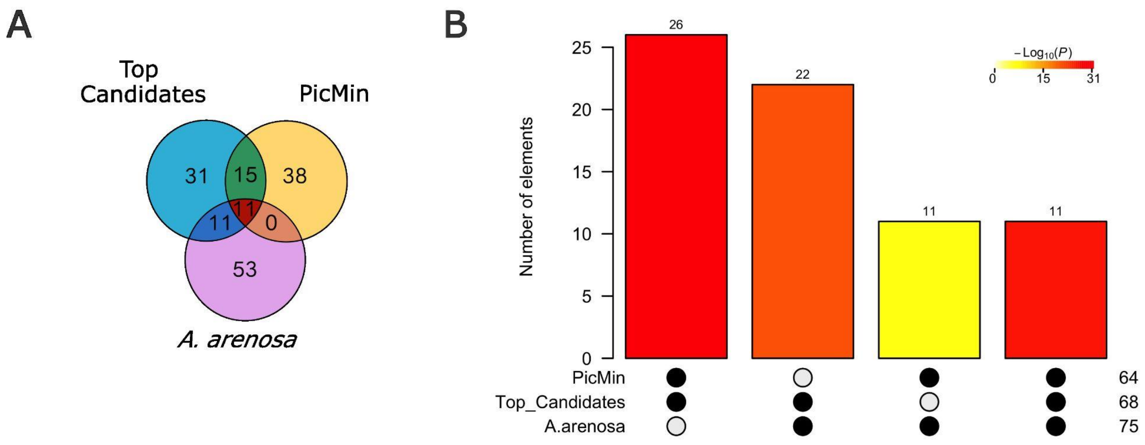


**Supplementary figure S14.** (A) Number of significantly enriched GO-terms overlapping between *Alyssum* candidate genes (top candidates and PicMin candidates) and *A. arenosa* candidate genes. (B) Significance of the intersection shown in A, tested using one-sided Fisher's exact test. Each intersection shows a higher than expected by chance only number of GO-terms overlapping ($p < 2.954653e-08$). The colour intensity of the bars represents the p-value significance of the intersections, and the number of elements shared is reported on each intersection bar.

| Methods | Metrics | Initial primary assembly | Final assembly |
|---|---|---|---|
| QUAST | Number of contigs | 874 | 168 |
| | Total length | 714798279 | 683338063 |
| | GC (%) | 38.51 | 38.45 |
| | N50 | 11862547 | 12254332 |
| | L50 | 18 | 17 |
| BUSCO | Complete (%) | 96.2 | 96.2 |
| | Single-copy (%) | 87 | 87 |
| | Duplicate (%) | 9.2 | 9.2 |
| | Fragmented (%) | 0.4 | 0.4 |
| | Missing (%) | 3.4 | 3.4 |
| Merqury | Quality value (QV) | 58.49 | 65.74 |
| | Completeness | 80.46 | 80.37 |

**Supplementary table S1:** Summary of genome assembly quality metrics assessed by BUSCO (completeness), QUAST (contiguity and assembly statistics), and Merqury (base-level accuracy and consensus quality values)
