## Supplementary Methods for "Genomic basis of adaptation to serpentine soil in two *Alyssum* species shows convergence with *Arabidopsis* across 20 million years of divergence"

**Determination of Population-specific Soil:**

- **Reaction-active and exchangeable:**

2,5 ml of homogenous soil sample (sieved through laboratory sieve with mesh grid unit diameter of 2 mm) was extracted for 30 minutes with use of orbital shaker. For this purpose, 25 ml of preboiled deionized water (active) or 0,1M KCl solution was added. After shaking soil suspensions were left to sit for another 90 minutes. Afterwards soil reaction was measured directly in suspensions by WTW Multilab 9620 IDS with combined pH electrode.

ISO 10390: Soil quality – Determination of pH, International Organization for Standardization, ISO 2000

- **Conductivity:**

10 g of homogenous soil sample (sieved through laboratory sieve with mesh mesh grid unit diameter of 2 mm) was weighed in to 250 ml extraction vessel, then 50 ml deionized water with specific conductivity ≤0,2 mS/m. The whole soil suspension is then extracted 30 minutes with use of orbital shaker. Afterwards the suspensions are filtered through thick filtration paper. For preparation of reaction blank same method is applied. Ideal value of blank is beneath 1 mS/m. Conductivity is then measured by WTW Multilab 9620 IDS with conductivity cell with thermo correction and automatic checking of conductivity constant.

ISO 10390: Soil quality – Determination of the specific electrical conductivity, International Organization for Standardization, 1992

- **Gravimetrical Determination of Dry mass and Water Content:**

5-10 g of air-dried homogenous soil sample (sieved through laboratory sieve with mesh grid unit diameter of 2 mm) is carefully weighed into the glass vessel and placed into the dried oven pre-heated up to 105°C for at least 6 h. After this time the glass vessels are tightly closed and leave to cool in laboratory desiccator for at least 30 minutes and is precisely weighed with accuracy of 0,001 g.

ISO/DIS 11465: Soil quality – Determination of the Dry Mass and Water Content on Mass Basis-Gravimetric Method, International Organization for Standardization, 1993

- **Exchangeable Phosphorus (Mehlich III Extract Solution):**

Homogenous soil sample (soil particles of size ≤ 2 mm) is extracted by extract solution Me III, in ratio of 5 g of air-dried soil to 50 ml of extract solution or 10 g of sample to 100 ml of extract solution, with use of orbital shaker. Afterwards the soil suspension is filtered through thick filtration paper and the clear filtrate is than further analysed. The amount of exchangeable phosphorus is determined spectrophotometrically as reaction product of direct reaction of phosphorus with ammonium molybdate with use of reaction mixture of sulphuric acid, ascorbic acid and potassium antimony tartrate. The colour of reaction product is blue, and the intensity of this coloration is determined by UV/Vis Unicam Spectrophotometer at wavelength of 750 nm.

- **Available Nutrients in Mehlich III Extract Solution:**

Air-dried soil samples homogenized on particle size ≤ 2 mm were extracted by group extract solution Me III, which contains ammonium fluoride, ammonium nitrate and ethylenediaminetetraacetic acid for support of desorption of exchangeable cations. Acidic reaction of extract solution is fixed by addition of acetic and nitric acid. For the extraction the rotary shaker was used. Afterwards the soil suspension is filtered through thick filtration paper and the clear filtrate is than further analysed. The content of available calcium and magnesium was determined by method of flame atomic absorption with addition of sulphuric acid and lanthanum chloride for elimination of phosphate and sulphate.

The content of other elements was determined by the same method just in original filtrate without further additions. Our laboratory uses AAS Spectrometer ContrAA 700 of Analytik Jena which for measurement of calcium, aluminium and chromium uses mixture of $N_2O$-$C_2H_2$ gases. For measuring of the other elements mixture of $C_2H_2$ with air was used.

- **Cation-exchange capacity:**

For purpose of this method 10 g of air-dried homogenous sample is weighed, then 100 ml of 0,1M $BaCl_2$ is added. The extraction vessels are closed and left to sit for 18 h. Afterwards soil suspensions are extracted by rotary shaker for 2 h and filtered through thick filtration paper. For the purpose of determination of calcium and magnesium the clear filtrate is diluted, and lanthanum chloride is added. Samples prepared by this method is then analysed by the method of atomic absorption. The content of sodium and potassium is determined at the original filtrate without further dilution nor addition. The analyses are then carried out by Spectrometer ContrAA 700 of Analytik Jena with use of mixture of $N_2O$-$C_2H_2$ (Ca) and $C_2H_2$ -air respectively. The content of hydrogen and aluminium ions is determined by titration of filtrate by 0,02M NaOH solution. The result is given at mmol chemical equivalent for kg of soil.

**<u>Determination of Ca, Mg, Co, Cr and Ni in plant tissues and soils relative to the transplant experiment</u>**

Determination of Ca, Mg, Co, Cr and Ni in plant tissues and soils used for the transplant experiment was carried using inductively coupled plasma optical emission spectrometry (ICP OES).

Due to very small amounts of plant samples in units to tens of mg, the samples were decomposed prior to the analysis using the microwave oven Speedwave ®Xpert (Berghof, Germany, maximal applied power 2000 W) with a multi-tube system. The plant tissue (2 replicates, 8 – 50 mg according to the available sample amount) was inserted into digestion tubes and treated with 2 ml of subboilingly destilled (Berghof, Germany) nitric acid (per analysis, Lachner, the Czech Republic) under the following conditions: 10 min hold on 170 °C,

30 % of maximal power, 10 min on 200 °C, 30 % of power, 30 min on 30 °C, 0 % of power. The mineralised samples were filled up to the final volume of 10 ml with deionised water (conductivity 0.055 μS/cm, Evoqua Water Technologies, Germany).

The elemental analysis of Ca, Mg, Co, Cr and Ni was carried out using the sequential, radially viewed ICP OES spectrometer INTEGRA 6000 (GBC, Dandenong Australia) equipped with the ultrasonic nebulizer U5000AT+ (Teledyne Cetac Technologies, the USA), concentric nebulizer (2 $ml.min^{-1}$) and a glass cyclonic spray chamber (both Glass Expansion, Australia). The analytical lines used were: Mg 285.2213 nm, Ca 422.673 nm, Ni 221.647 nm, Co 238.892 nm and Cr 267.716 nm. The operation conditions of the ICP OES analysis were as follows: sample flow rate 1.5 $mL.min^{-1}$, plasma power 1000 W, plasma, auxiliary and nebulizer gas flow rates 10, 0.4, and 0.52 $L.min^{-1}$, respectively, photomultiplier voltage 600 V for Ni, Co and Cr and 350 V for Ca and Mg, view height 6.5mm, three replicated reading on-peak 1 s, fixed point background correction. The multielemental standards containing 10 – 5 – 1 – 0.5 – 0.1 $mg.L^{-1}$ of Mg and Ca and 0.1 – 0.05 – 0.01 – 0.005 – 0.001 $mg.L^{-1}$ Ni, co and Cr were used for instrument calibration. The external calibration standards were prepared using standard solutions of Mg, Ca, Ni, Co and Cr all containing 1 $g.L^{-1}$ (SCP, Canada). The limits of detection (concentration equal to three times the standard deviation at the point of the background correction) were 0.0005 $\mu g.L^{-1}$ for Ni, Co and Cr and 2 $\mu g.L^{-1}$ for Mg and Ca. Certified reference material (Bush twigs and leaves GBW 07602 from the China National Analysis Center for Iron and Steel, Beijing) was used to validate the method and for the quality control.

Analysis of soils was carried out using the Mehlich 3 extraction protocol (see link below) followed by ICP OES method as described for plants.0.5 g of sample was treated with 50 ml of the Mehlich 3 extraction solution for 10 min using a laboratory shaker, then 10 min left to settle, then filtered and and appropriately diluted prior to analysis.
